# Variation in the fitness impact of translationally optimal codons among animals

**DOI:** 10.1101/2024.07.22.604600

**Authors:** Florian Bénitìere, Tristan Lefébure, Laurent Duret

## Abstract

Early studies in invertebrate model organisms (fruit flies, nematodes) showed that their synonymous codon usage is under selective pressure to optimize translation efficiency in highly expressed genes (a process called translational selection). In contrast, mammals show little evidence of selection for translationally optimal codons. To understand this difference, we examined the use of synonymous codons in 223 metazoan species, covering a wide range of animal clades. For each species, we predicted the set of optimal codons based on the pool of tRNA genes present in its genome, and we analyzed how the frequency of optimal codons correlates with gene expression to quantify the intensity of translational selection (*S*). Surprisingly, few metazoans show clear signs of translational selection. As predicted by the nearly neutral theory, the highest values of *S* are observed in species with large effective population sizes (*N* _e_). Overall, however, *N* _e_ appears to be a poor predictor of the intensity of translational selection, suggesting important differences in the fitness effect of synonymous codon usage across taxa. We propose that the few animal taxa that are clearly affected by translational selection correspond to organisms with strong constraints for a very rapid growth rate.

## Introduction

Since the early days of DNA sequencing, it has been noticed that the usage of synonymous codons is not random: some synonymous codons are more frequently used than others, and the patterns of synonymous codon usage (SCU) can vary both across species and among genes within a genome (Grantham *et al*., 1980). Two types of processes, adaptive or non-adaptive, can contribute to genome-wide patterns of SCU (Sharp *et al*., 1993). First, neutral substitution patterns (NSPs) vary across taxa and, in some species, can also vary along chromosomes. NSPs are primarily driven by the underlying pattern of mutation, which accounts for 60% of the variance in genome base composition across the tree of life (Long *et al*., 2018). In addition, in some taxa, NSPs are also strongly affected by GC-biased gene conversion (gBGC), a process associated to meiotic recombination that favors the transmission of G:C alleles over A:T alleles (Duret and Galtier, 2009). NSPs affect all genomic compartments (coding or non-coding), and notably have a strong impact on SCU (*e.g.* Pouyet *et al*. (2017); Long *et al*. (2018)). Besides NSPs, SCU can also be affected by selection. Indeed, it has been observed that in some species, SCU varies according to gene expression level (Gouy and Gautier, 1982; Sharp *et al*., 1986; Duret and Mouchiroud, 1999) and that the synonymous codons that are more frequently used in highly expressed genes correspond to the most abundant tRNAs (Ikemura, 1985; Dong *et al*., 1996; Moriyama and Powell, 1997; Kanaya *et al*., 1999; Duret, 2000). This indicates that synonymous codon usage and tRNA content have coevolved in a way that optimizes translation. This co-evolution implies two types of selective pressure (Bulmer, 1987): 1) selection on the pool of tRNAs to match the relative abundance of different codons in the transcriptome (*i.e.* the codon demand), and 2) selection on the synonymous codon usage of genes to match the pool of tRNAs, referred to as “translational selection”.

It is generally considered that there are two main benefits of using translationally optimal codons. First, this leads to faster translation, and hence to reduce the time spent by ribosomes on each mRNA, thereby increasing the pool of free ribosomes available in the cell, which ultimately allows a higher cellular growth rate (Bulmer, 1991). Second, the usage of synonymous codons decoded by the most abundant tRNAs increases the accuracy of translation, and thus reduces the amount of mis-translated proteins that cause an important burden on the cell (Akashi, 1994; Drummond *et al*., 2006). It is important to note that for both aspects (speed and accuracy of translation), the benefit of using optimal codons is expected to be proportional to gene expression level. First, the higher the expression level of a given gene, the stronger the impact of its translation speed on the pool of free ribosome. Second, for a given mis-translation rate, the cost of erroneous protein production (in terms of waste of resources and of direct toxic effect of misfolded proteins) increases directly with expression level. In bacteria, the intensity of translational selection is correlated to the minimal cell division time, which suggests that the selective force for the optimization of SCU is the maximization of cellular growth (Rocha, 2004; Sharp *et al*., 2005).

It should be noted that besides translational selection, synonymous sites can be subject to additional levels of selective constraints. For instance, the presence of splice enhancers located within exons skews codon usage near exon-intron boundaries in mammalian genes (Parmley and Hurst, 2007). However, this type of selective pressure is site-specific (*i.e.* a particular codon is preferred at a specific site in a given gene), and hence, is not expected to affect the genome-wide pattern of SCU. Similarly, there is evidence that the use of translationally sub-optimal codons can be advantageous at some specific sites to slow-down translation and favor the proper folding of proteins (Buhr *et al*., 2016; Walsh *et al*., 2020). But again, this is expected to be a local effect, with limited genome-wide impact on SCU.

Interestingly, the intensity of translational selection varies widely across species, not only in unicellular organisms, but also in multicellular eukaryotes (Sharp *et al*., 2005; Subramanian, 2008; dos Reis and Wernisch, 2009; Galtier *et al*., 2018). For instance, among animals, early studies on the two main invertebrate model organisms (the fruitfly *Drosophila melanogaster* and the nematode *Caenorhabditis elegans*) showed clear signatures of translational selection (Shields *et al*., 1988; Duret and Mouchiroud, 1999), whereas there is no sign of translational selection in humans (Śemon *et al*., 2006; Pouyet *et al*., 2017). It should be emphasized that there is clear evidence for an effect of SCU on gene expression in mammals (Kudla *et al*., 2006; Courel *et al*., 2019; Wu *et al*., 2019; Mordstein *et al*., 2020; Medina-Muñoz *et al*., 2021), and in practice, codon optimization is essential for the design of heterologous gene expression systems, notably for human mRNA vaccines (Leppek *et al*., 2022). So if SCU affects gene expression in humans, why is this trait not under selective pressure? To address this point, and more generally to understand variation in the intensity in translational selection across animals, it is important to refer to population genetics principles (Ohta, 1996). Indeed, the SCU in a given genome reflects a balance between selection favoring translationally optimal codons, and the effects of mutation and drift, allowing the fixation of non-optimal codons (Bulmer, 1991). Thus, the frequency of optimal codons is expected to depend on the mutational pattern and on the population-scaled selection coefficient (*S* = 4*N*_e_*s*), where *N* _e_ is the effective population size and *s* the selection coefficient in favor of translationally optimal codons (Bulmer, 1991; Sharp *et al*., 2005). Hence, the lack of translational selection in some animal taxa might stem from a small *N* _e_ (hereafter referred to as the drift-barrier hypothesis), or from a smaller fitness effect of using translationally optimal codons (*i.e.* lower *s*).

To explore these hypotheses, several previous studies analyzed variation in the intensity of translational selection across eukaryotes (Subramanian, 2008; dos Reis and Wernisch, 2009; Galtier *et al*., 2018). These three studies, reported positive correlations between signatures of translational selection and proxies of *N* _e_. Although this pattern is qualitatively consistent with the predictions of the drift barrier hypothesis, quantitatively, the observations do not fit with this model. For instance, dos Reis and Wernisch (2009) estimated *S* in 10 eukaryotic species, and they reported only a 2-fold difference in *S* between humans and *D. melanogaster* (respectively *S* = 0.5 and *S* = 1.0), despite a *≈*30-fold difference in *N* _e_ between the two species (20,000 vs. 600,000; Lynch *et al*. (2023)). According to the authors, this poor fit to the drift barrier model might be due to the fact that their analysis was sensitive to variation in NSP across genes, which might have led to overestimate *S* in humans (dos Reis and Wernisch, 2009). But, it has also been argued that besides differences in *N* _e_, *s* is also likely to vary across species, as long-lived organisms, with relatively slow development, are likely to be less constrained to optimize cell growth than species with a very rapid development (Subramanian, 2008).

These three studies were based on relatively limited sample sizes (10 to 30 species), and in the end, the causes of the variation in the intensity of translational selection across species remained unclear. To try to go further, we decided here to investigate variation in translational selection intensity across a large dataset of 223 metazoan species, covering a wide range of animal clades. For each species, we predicted the set of optimal codons based on the pool of tRNA genes present in its genome, and we analyzed how the frequency of optimal codons varies with gene expression, controlling for variation in NSP. Based on these variations, we quantified *S* in each species, and analyzed how it correlates with estimates of *N* _e_ or life history traits. Our analyses revealed that overall, few metazoans show clear signs of translational selection. As expected, the highest values of *S* are observed in species with large *N* _e_, while species with small *N* _e_ show little evidence of translational selection. However, overall, *N* _e_ appears to be a poor predictor of the intensity of translational selection, which suggests important variation in *s* across taxa. We discuss factors that may drive this variation in the fitness effect of optimizing codon usage.

## Results

### Non-adaptive processes are the primary drivers of codon usage variation among metazoans

To investigate the factors driving the intensity of translational selection in metazoans, we used the GTDrift database, that compiles genomic and transcriptomic data along with life history traits and proxies of *N* _e_ for various eukaryotic species (Bénitìere *et al*., 2024). We initially selected 257 metazoan species available in GTDrift, but we excluded 11 species for which there were not enough transcriptomic data to quantify the distribution of expression level (less than 5,000 genes detected as being expressed). We analyzed patterns of SCU and genomic base composition in the 246 remaining species, covering a wide range of clades (129 vertebrates, 82 insects and 35 other metazoan species; Fig. 1A).

**Figure 1:**
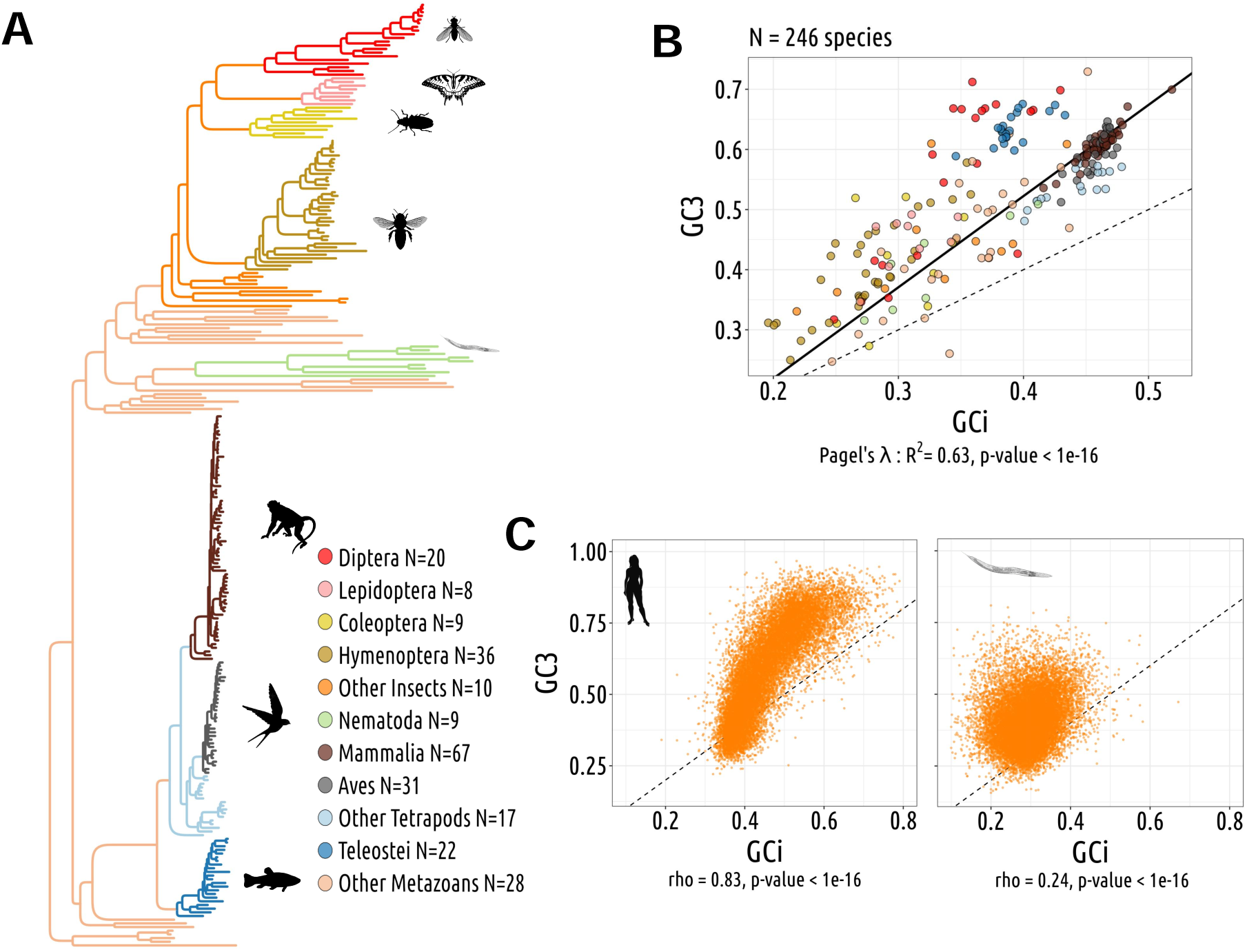
Codon usage variations are driven by non-adaptive processes. **A**: Phylogenetic tree of the 257 studied species. **B**: Gene average GC content at the third position of codons (GC3) and the gene average GC in introns (GCi) for each species. Pagel’s *lambda* model is used to take into account the phylogenetic structure of the data in a regression model (black line). **C**: Correlation between GC3 and GCi in *Homo sapiens* (left) and in *Caenorhabditis elegans* (right), each point corresponds to one gene. Spearman’s rho and corresponding p-values are displayed under the graph. The dotted lines correspond to x=y.

Patterns of SCU can be affected both by translational selection and by NSPs (Sharp *et al*., 1993). It is possible to distinguish the contribution of NSPs because they affect the base composition of both coding and non-coding regions, whereas translational selection operates only on codons. Thus, if differences in SCU across species are driven by NSPs, then it is expected that they should be correlated with variation in the base composition of non-coding regions. Similarly, if intra-genomic variation of SCU in a given species is driven by the heterogeneity of NSPs along its chromosomes, then this should result in a covariation between the codon usage of genes and the base composition of their introns. Owing to the symmetry of the DNA molecule, NSPs generally affect similarly both strands, resulting in an equal proportion of cytosine (C) and guanine (G), as well as an equal proportion of thymine (T) and adenine (A) (Lobry, 1995). Hence, the G+C content provides a good summary statistics of the impact of NSPs on the genomic base composition. To examine the potential contribution of non-adaptive processes to the observed variations in SCU across the 246 species, we measured their G+C content in introns (GCi) and at the third position of codons (GC3), averaged over all genes. We observed a strong correlation between the average GC3 and the average GCi (Fig. 1B). This indicates that non-adaptive processes, are the primary factor driving the observed variation in codon usage across species.

NSPs can vary within the genome of a given species, and impact codon usage accordingly. In *Homo sapiens*, the *per* gene GC3 and GCi are highly correlated (Spearman’s correlation coefficient, rho=0.83, p<10^−16^), whereas this correlation is less pronounced in *Caenorhabditis elegans* (rho=0.24, p<10^−16^; Fig. 1C). Species showing the strongest intra-genomic variance in codon usage (as assessed by GC3) are the ones with the strongest variance in GCi Supplementary Fig. 1A). These correlations between GC3 and GCi are particularly strong in tetrapods (107/108 species with rho>0.7) and in hymenopterans (25/35 species with rho>0.7) (Supplementary Fig. 1B). The other clades generally show less variance in GCi, and weaker correlations between GC3 and GCi. But overall, 246/246 species (100%) showed a significant positive correlation (p<0.05), which indicates that in most species, intra-genomic variation in NSPs somehow contribute to the variance in SCU among genes. Hence, it is important to take this source of variance into account to be able to detect signatures of translational selection within genomes.

### tRNA abundance matches proteome requirements

To quantify the intensity of translational selection, we used an approach similar to that of dos Reis and Wernisch (2009) and Sharp *et al*. (2005). This approach is based on the comparison of the frequency of optimal codons between highly and weakly expressed genes, and therefore requires the prior identification of optimal codons. For this, dos Reis and Wernisch focused on the nine amino acids that are encoded by two codons (duet codons), and predicted the optimal codon of each amino acid as being the one that is more frequently used in highly expressed genes. One caveat is that if the NSP varies among genes according to their expression level, this may lead to erroneous prediction of codon optimality. Furthermore, this approach does not capture the signal of translational selection from the nine other amino acids that are encoded by triplet, quartet or sextet codons. To avoid these limitations, we sought here to predict optimal codons based on the tRNA pool. Owing to technical difficulties, there are currently few species for which tRNA abundance has been quantified directly. Behrens *et al*. (2021) recently developed a technique (mim-tRNAseq) that allowed them to measure tRNA abundance in four eukaryotes (Behrens *et al*., 2021). This study revealed a robust correlation between tRNA abundance and their respective gene copy number, with an adjusted *R*^2^ > 0.91 for yeasts (*S. cerevisiae* and *S. pombe*), 0.79 for *Drosophila melanogaster* and 0.62 for *Homo sapiens* (Behrens *et al*., 2021). These results suggest that tRNA gene copy number is a good predictor of tRNA abundance.

To investigate whether the number of tRNA genes could be used as an indirect measure of tRNA abundance across metazoans, we analyzed the co-variation of their tRNA gene repertoires with the amino acid composition of their proteome. The total number of tRNA gene copies varies widely among clades and species (ranging from an average of 201 tRNA gene copies per genome in hymenopterans to 1,539 copies in teleost fish; Supplementary Fig. 2). However, the relative copy number of distinct isoacceptor tRNA genes is quite conserved among metazoans (Supplementary Fig. 3). There are some rare cases where the gene copy number of a given tRNA has exploded in a given species compared to other genomes (Supplementary Fig. 2). This might reflect the propensity of tRNA genes to become transposable elements (for instance, the genome of *Blattella germanica* contains 711 copies of the AGA Ser-tRNA, vs. 1 to 128 copies for the other tRNAs). Indeed, many SINE retrotransposon families derive from tRNA genes (Sun *et al*., 2007), and it is therefore possible that some recently evolved SINEs are erroneously annotated as *bona fide* tRNA genes.

In both *Drosophila melanogaster* and *Homo sapiens*, we observed a strong correlation between amino acid usage (*i.e.* the frequency of amino acids, weighted by the gene expression level) and direct measures of tRNA abundance (rho=0.79; Supplementary Fig. 4A,B; Behrens *et al*. (2021)). These results indicate that tRNA abundance matches the amino acid demand. As expected, the amino acid usage of these two species also strongly correlates with their tRNA gene copy numbers (rho=0.78 and 0.68 respectively; Supplementary Fig. 4C,D). To assess the generality of this relationship, we computed the correlation between tRNA gene copy number and the amino acid demand across 246 animal species. We observed a significantly positive Spearmann coefficient (p-value<0.05) in 93% of species (Fig. 2). This indicates that in most of metazoans, the tRNA gene copy number is under constraints to match the amino acid demand. This implies that tRNA abundance is primarily regulated by modulating the copy number of tRNA genes rather than their transcription level. We suspect that the few cases where the number of tRNA genes does not correlate with amino acid usage might be due to annotation errors: some tRNA genes may have been missed (*e.g.* because of gaps in the genome assembly), or conversely, some SINEs or tRNA pseudogenes may have been incorrectly annotated as functional tRNA genes. To ensure that the tRNA gene copy number is a good proxy of the tRNA abundance, we kept in our study only the species for which tRNA gene copy number correlates significantly with amino acid usage (N=230 species). We also excluded 7 species for which the repertoire of annotated tRNA appeared to be incomplete (*i.e.* the cognate tRNAs of certain codons were not found in the genome assembly).

**Figure 2:**
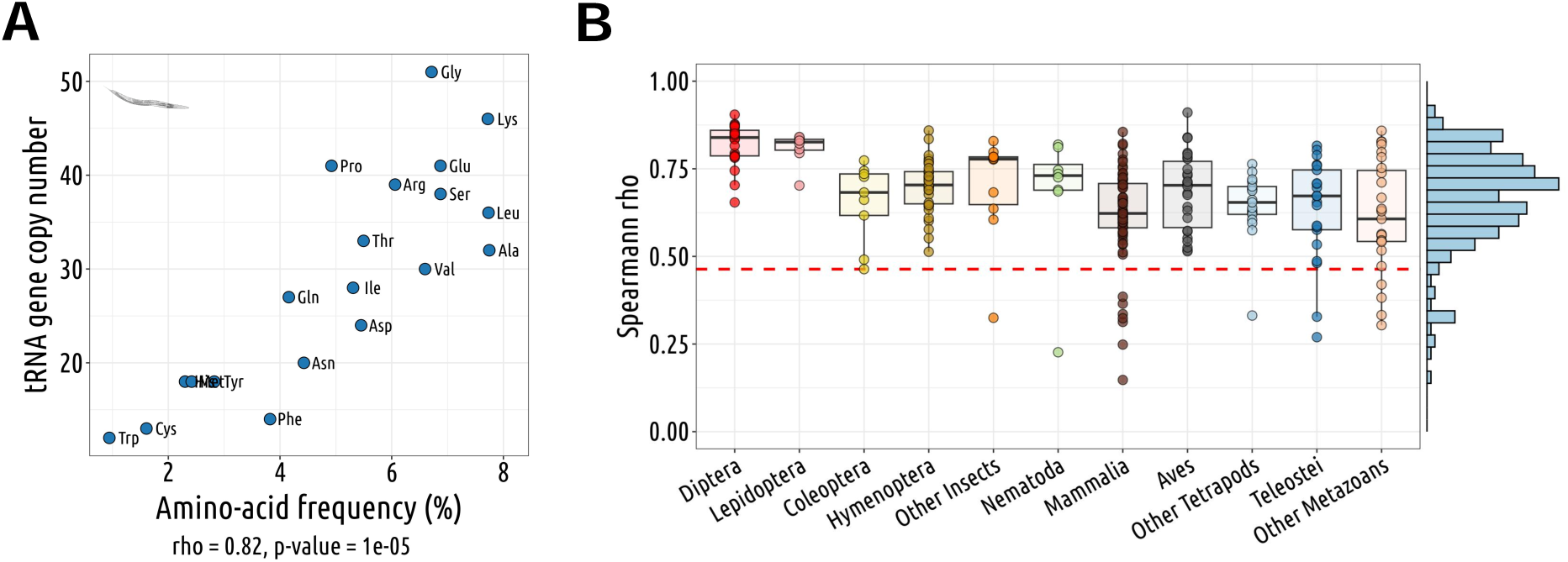
The tRNA gene copy number is a good predictor of the codon demand. **A**: Representative example of the relationship between the number of isoacceptor tRNA gene copy *per* amino acid and the frequency of amino acid weighted by gene expression in *Caenorhabditis elegans*. Spearman’s rho and corresponding p-value are displayed under the graph. **B**: Boxplot showing the distribution of Spearman correlation coefficients (ρ) from Panel A for each species (N=246 species). The red line indicates the threshold above which the p-value is lower than 0.05.

### Definition of putative-optimal codons based on tRNA abundance and wobble-pairing rules

To predict which synonymous codons are optimal for translation, it is first necessary to associate each of the 61 codons to their cognate tRNA. The number of distinct isodecoder tRNAs (*i.e.* distinct anticodons) ranges from 43 to 60 per species (average=47). This implies that 1 to 18 codons cannot be translated through Watson-Crick pairing (WCp), and hence have to be translated via wobble pairing (WBp). We used the rules established by Percudani (Percudani, 2001) to assign each of these codons to their cognate tRNA, allowing for non-standard base pairing with the first nucleotide of the anticodon (Fig. 3A). For example, deamination of adenine in inosine (I) in anticodons ANN makes them permissive to wobble pairing I:C; I:U or I:A. Another common wobble pairing is the G:U/U:G pairing (Percudani, 2001). As an illustration, in human, asparagine is translated by a single tRNA (anticodon GTT) that decodes both AAC (by WCp) and AAT (by G:U WBp). AAT accounts for 48% of asparagine codons, highlighting the significance of wobble pairing.

**Figure 3:**
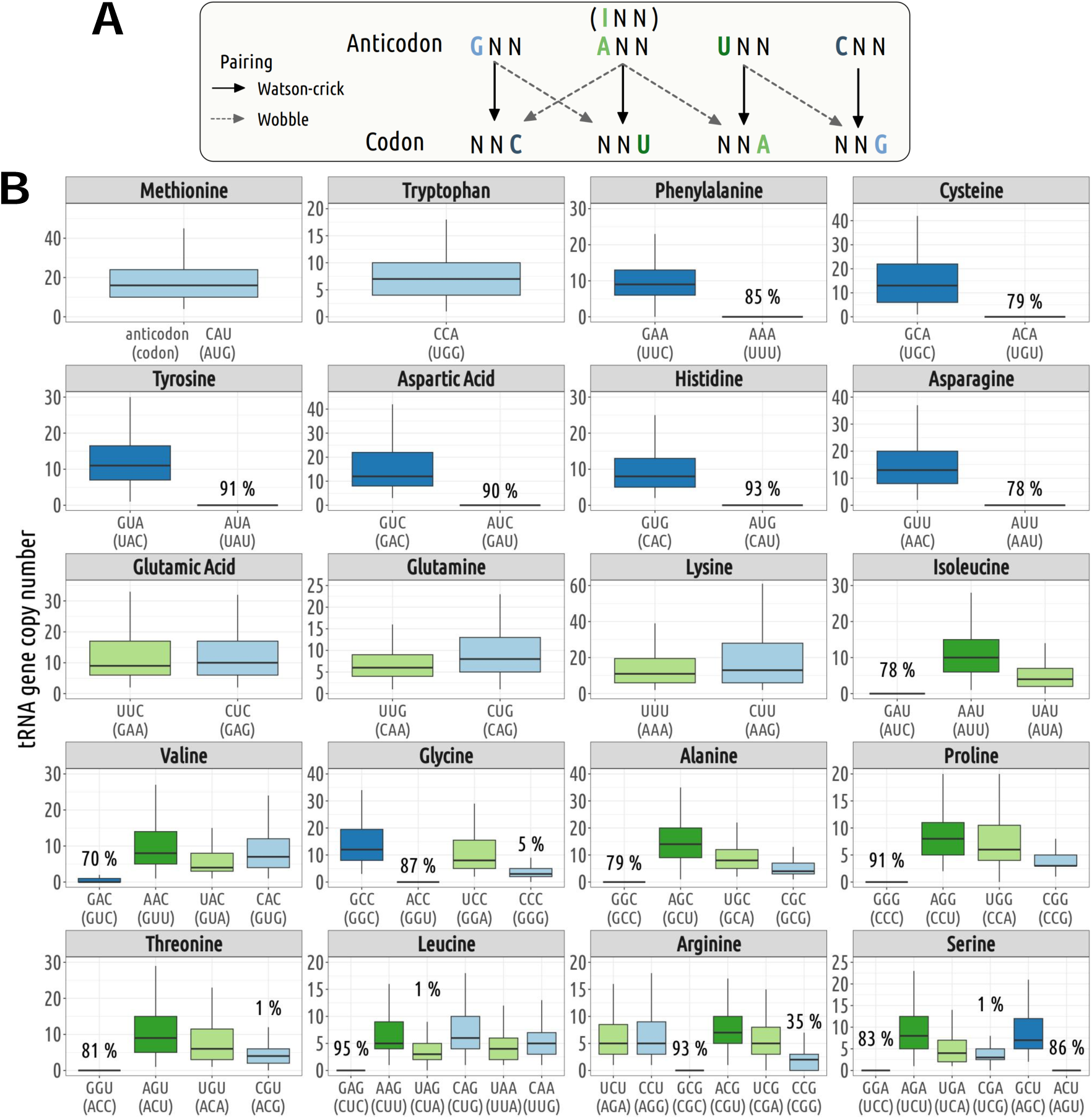
Codon-anticodon assignment. **I**n each genome, we used the rules proposed by Percudani (Percudani, 2001) to assign codons to their cognate tRNA: first, isodecoder tRNAs present in a given genome are assigned to their complementary (Watson-Crick) codons; then, the remaining codons (for which the genome does not contain any tRNA carrying the complementary anticodon), are inferred to be decoded by wobble pairing. A: Illustration of the various possible pairings: Watson-Crick and wobble pairing. **B**: Distribution of tRNA gene copy number across 223 species. The boxplot represents the median, interquartile range (box edges at 25th and 75th percentiles), and whiskers extending to the largest value no further than 1.5 times the interquartile range. For tRNA isodecoders that are absent from some genomes, the percentage of species where they are missing is also indicated.

There are 18 amino acids that are encoded by multiple synonymous codons. These amino acids can be classified in two groups: (i) those whose synonymous codons are translated by at least two distinct isodecoder tRNAs, and (ii) those for which all synonymous codons are translated by a single isodecoder tRNA. There is some variation in the set of amino acids present in each group, depending on the isodecoder tRNA repertoire of each species. The first group generally corresponds to amino acids encoded by sextet codons (Leu, Arg, Ser), quartets (Val, Gly, Ala, Pro, Thr), triplet (Ile) and NNG/NNA duets (Glu, Gln, Lys). The second group corresponds essentially to the six amino acids encoded by NNC/NNT duets (Phe, Cys, Tyr, Asp, His, Asn) (Fig. 3B).

In the following analyses, for each amino acid of the first set, synonymous codons were predicted to be optimal if they were decoded by the isodecoder tRNA with the highest gene copy number (*i.e.* predicted to be the most abundant). In case of ex æquo (*i.e.* if all synonymous codons are decoded by isodecoders having the same gene copy number), then the optimal codons of this amino acid were considered as unknown (76 ex æquo among 785 cases in total, in 62 species). In the cases where the most abundant tRNA decodes more than one synonymous codon, we considered all of them as potentially optimal (*i.e.* at this stage, we do not make any assumption regarding which of the Watson-Crick pairing or wobble pairing is the most efficient). This first set of putative-optimal codons will hereafter be referred as “POC1”. The second set of amino acids corresponds to cases where the two synonymous codons (NNC/NNT) are decoded by a single isodecoder (anticodon GNN). There is evidence, based on studies in various eukaryotes, that the wobble pairing GNN:NNU is less efficient than the Watson-Crick pairing GNN:NNC (Stadler and Fire, 2011; Chan *et al*., 2017; Wang *et al*., 2017). Consequently, for these amino acids, we defined codons NNC (decoded through WCp) as being the putative-optimal codons “POC2”.

For the human genome, POC1 can be defined for 13 amino acids and POC2 for 5 amino acids. For *Caenorhabditis elegans*, POC1 and POC2 are defined for 12 and 6 amino acids, respectively. On average among the 223 species, POC1 are defined for 12.5 amino acids *per* species (ranging from 8 to 17) and POC2 for 5.2 amino acids (ranging from 0 to 6, except *Tyto alba* with 7 POC2, including Ile).

### Highly expressed genes are enriched in optimal codons

The intensity of translational selection depends directly on gene expression levels. Given the very wide range of variation of gene expression levels (> 1000 folds), the fitness impact of synonymous codon usage is expected to vary strongly among genes. Hence, a typical feature of genomes subject to translational selection (TS), is that the frequency of optimal codons is particularly high in the most highly expressed genes. Thus, to identify which species are subject to TS, we examined the variations in POC frequency according to gene expression level. To control for possible variations in neutral substitution patterns, we also analyzed the corresponding triplet usage in introns, referred to as POC-control. It is important to note that even in absence of TS, the frequency of POC-control is not necessarily expected to be equivalent to that of the overall POC frequency, notably because the base composition of non-coding regions is affected by indels and transposable elements, which are counter-selected in coding regions.

For *Homo sapiens* POC frequencies show some slight fluctuations with gene expression (Fig. 4A). However, the same weak fluctuations are observed for POC-controls in introns, which implies that the same process, independent of translation efficiency, affects the base composition, both in introns and at synonymous codon positions (Fig. 4A). Indeed, there exists a strong correlation between the GC content of introns and the GC3 content in *Homo sapiens*, along with pronounced variations in GC3 content (Fig. 1C). In contrast, in *Caenorhabditis elegans*, we observed a strong rise in POC frequencies in highly expressed genes, both for POC1 (from 47% to 76%) and for POC2 (from 38% to 70%). These changes in codon usage are not caused by shift in local substitution patterns as we see no similar variation in POC-control (Fig. 4B).

**Figure 4:**
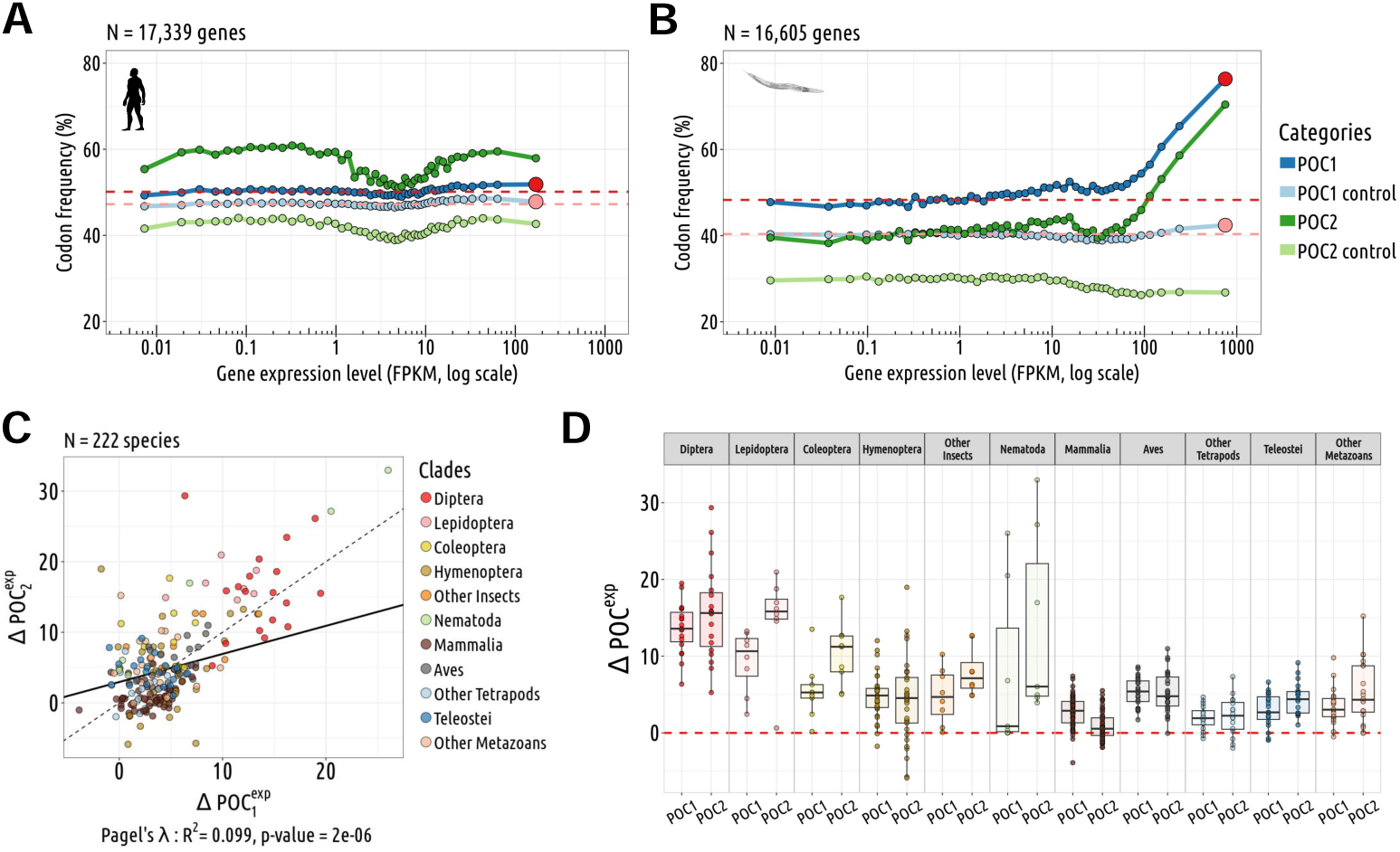
Relationship between the frequency of putative-optimal codons and gene expression level. **A,B**: Variation in the proportion of POC within coding sequences (POC1: dark blue; POC2: dark green) according to gene expression level. To control for variations in neutral substitution patterns, we analyzed the frequency of corresponding triplets within introns (POC1 control: light blue; POC2 control: light green). Each point represents a 2% bin of genes, with the red point at the end of each POC1 curve denoting the 2% most highly expressed genes. The red lines indicate the average POC1 frequency observed in the 50% least expressed genes (FPKM). The difference in POC frequency between the top 2% most highly expressed genes and the 50% least expressed genes is noted Δ*POC^exp^*. **A** represents *Homo sapiens*, and **B** represents *Caenorabditis elegans*. **C**: Correlation between 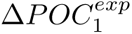 and 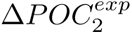 **D**: Δ*POC^exp^* distributions per species and by clades for the POC1 and POC2 (N=223 species).

It is important to notice that the non-linear relationship observed between gene expression level and POC frequency is perfectly consistent with the TS model, that assumes that the selection coefficient on synonymous codon usage (*s*) should increase linearly with gene expression level. Indeed, this model predicts that for lowly expressed genes (such that *S* = 4*N* _e_*s ≪* 1), the frequency of optimal codons should evolve neutrally, and hence should be independent of expression level. But above the ‘nearly-neutral’ point (*i.e.* the expression level for which *S ≈* 1), the frequency of optimal codon should strongly increase with expression level. The shape of the POC1 and POC2 curves in Fig. 4B indicates that in *C. elegans*, this ‘nearly-neutral’ point is reached for a gene expression level of about 50 FPKM. This implies that genes with a lower expression level (which represent 83% of genes in *C. elegans*) are not affected by TS. Of note, all else being equal, the fraction of genes affected by TS is expected to be even more reduced in species with a lower effective population size.

To assess the impact of TS on synonymous codon usage, we measured the difference between the frequency of POCs in the most expressed genes (top 2%), and the frequency of POCs in the 50% lowest expressed genes, controlling for POCs-control variations (see Materials & Methods). This shift in codon usage (denoted Δ*POC^exp^*) was computed for both POC1 and POC2 codons in each of the studied species (N=223 species). 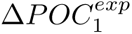 and 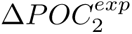 are strongly correlated (*R*^2^ = 46%, p-value*<10^−^*^16^), which indicates that the signature of translational selection is effectively captured by both sets of codons (Fig. 4C). For 211 species (95%), the frequency of POC1 is higher in highly expressed genes (Fig. 4D). Similarly, 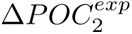 is positive for 191 species (86%; Fig. 4D). The higher Δ*POC^exp^* were observed in *C. elegans* (+30% for both POC1 and POC2). There are substantial variation across clades: average 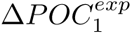 and 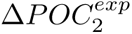 values are around +14% in Diptera compared to +3% in vertebrates. We obtained very similar results when measuring Δ*POC^exp^* without accounting for POC-control variations (Supplementary Fig. 5). Given that the two sets of codons gave very consistent results, we hereafter considered the whole set of POCs regrouping both POC1 and POC2.

### Weak relationship between the strength of translational selection and the effective population size

According to standard population genetic models of translational selection (Bulmer (1991); Sharp *et al*. (2005); dos Reis and Wernisch (2009)), the difference in codon usage between highly and weakly expressed genes is expected to be directly linked to the population-scaled selection coefficient in favor of optimal synonymous codons (*S* = 4*N*_e_*s*). Indeed, considering that synonymous codon usage evolves neutrally in lowly expressed genes, then *S* in highly expressed genes can be expressed as:

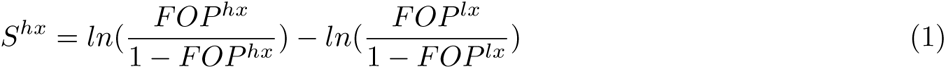

where *FOP ^hx^* and *FOP^lx^* are the observed frequencies of optimal codons in highly and lowly expressed genes respectively (Sharp *et al*., 2005; dos Reis and Wernisch, 2009). It should be noted however that this equation holds true only if underlying mutation patterns (and possibly gBGC) do not vary with gene expression level (Sharp *et al*., 2005; dos Reis and Wernisch, 2009). We used the above equation to estimate *S*^hx^ in each species, based on the observed POC frequencies in the top 2% most highly expressed genes, compared to the 50% least expressed. The choice of this latter threshold is based on the observation that in species with clear signature of translational selection, POC frequencies show little variation in genes below the median expression level (Fig. 4B; Supplementary Fig. 6).

If constraints on synonymous codon usage are similar across species (*i.e.* if *s^hx^* is constant), then *S^hx^* is expected to vary linearly with the effective population size (*S*^hx^ = 4*N*_e_*s*^hx^). To test this prediction, we sought to estimate *N* _e_ for each species. Lynch and colleagues recently compiled a list of species for which the germline mutation rate (µ) and the level of neutral diversity (*π_s_*) have been measured, and hence for which it is possible to infer the effective population size (*N_e_* = *π_s_*/4*µ*) (Lynch *et al*., 2023). This list included 24 species of our data set. We expanded this data set by including *N* _e_ estimates for 17 additional congeneric species. To explore the relationship between *S*^hx^ and *N* _e_ in more species, we also used three indirect proxies (longevity, body length and the *dN/dS* ratio) that correlate with the effective population size (Supplementary Fig. 7).

Among the 223 species analyzed, the strongest intensity of selection is observed in the nematodes *C. elegans* (*S*^hx^ = 1.3) and *C. nigoni* (*S*^hx^ = 1.0; Fig. 5). Dipters also show relatively strong values of *S*^hx^(mean = 0.63±0.16 sd), followed by lepidopters (mean *S*^hx^ = 0.41±0.15 sd). In vertebrates, signals of translational selection are weak (mean *S*^hx^= 0.15±0.12 sd), but nevertheless, *S*^hx^ are on average significantly non null (Student’s t-Test, p-value *<10^−^*^16^). As predicted by the drift barrier model, the species with the strongest signs of translational selection all show a relatively short lifespan, low body mass and low *dN/dS* (Fig. 5A,B and C), *i.e.* traits associated to organisms with large *N* _e_. Conversely species with traits associated to low *N* _e_ all show low *S*^hx^. However, the correlations between *S*^hx^ and *N* _e_ proxies are weak, and significant for only two of them (longevity and *dN/dS*) (Fig. 5A,C). The weakness of these correlation might be due to the fact that these traits are only indirect proxies of *N* _e_. However, even for the few species for which it is possible to get more direct estimates of *N* _e_ (based on *π_s_* and µ), the correlation between *S*^hx^ and *N* _e_ remains weak (Fig. 5D).

**Figure 5:**
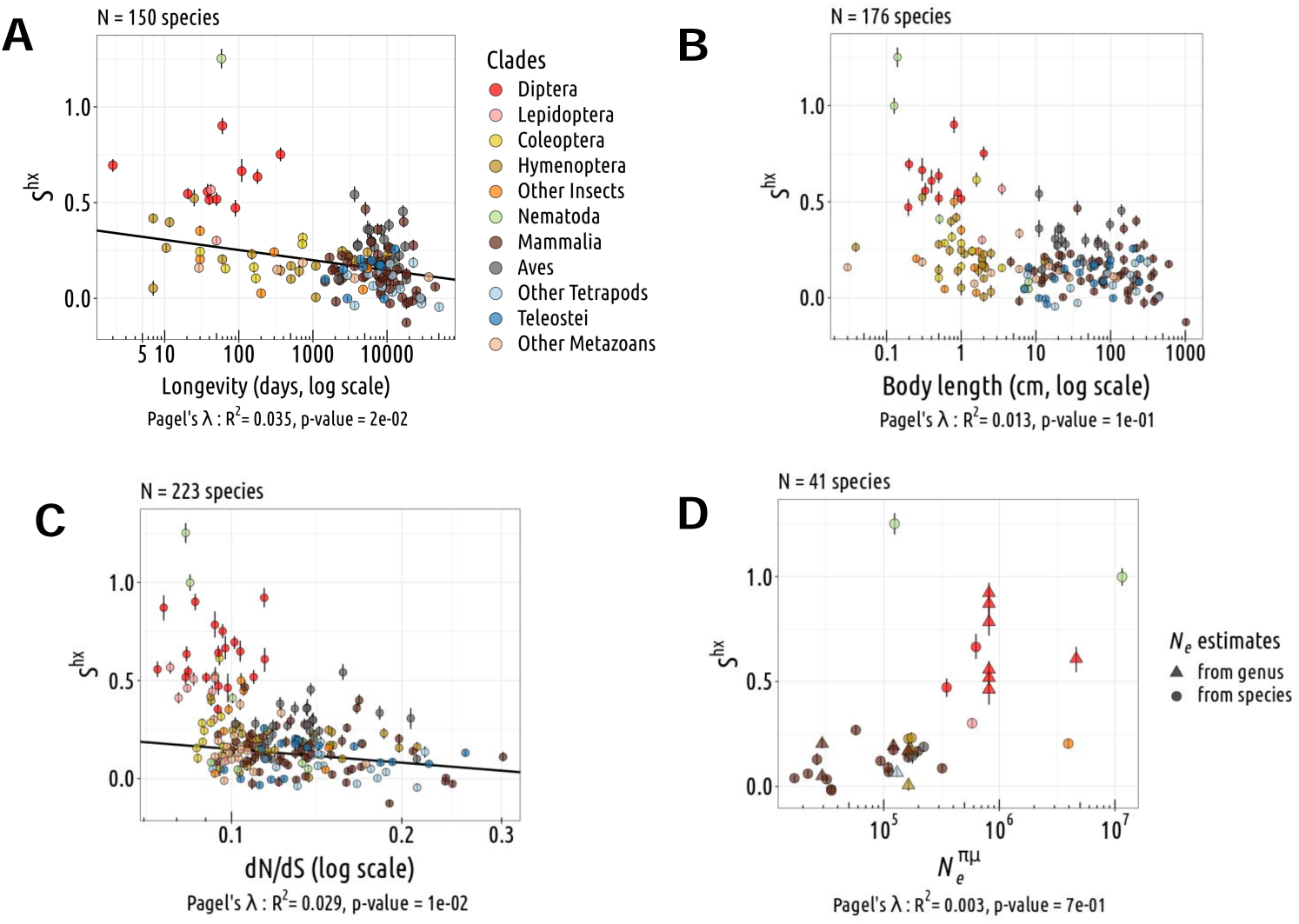
Relationship between *N* _e_ and translational selection intensity. Relationship between the population-scaled selection coefficient (*S*) and longevity (**A**), body length (**B**), *dN/dS* (**C**), *N ^πµ^* (**D**) (N=223 species). The translational selection intensity *S* is measured on the top 2% most highly expressed genes (*S*^hx^). Pagel’s *lambda* model is used to take into account the phylogenetic structure of the data in a regression model (the regression line is displayed in black when the correlation is significant). Error bars represent the 2.5th and 97.5th percentiles of *S*^hx^ values obtained by bootstraping (1000 draws with replacement among the top 2% most highly expressed genes, and the 50% least expressed).

### In species subject to translational selection, the tRNA pool evolves in response to changes in neutral substitution patterns

The above analyses show that for most metazoan species, translational selection is very weak, and hence that their synonymous codon usage is essentially shaped by neutral substitution patterns (NSP). Interestingly, even in species with clear signal of translational selection, codon usage appears to be influenced by variations in NSP. Notably, Diptera and Lepidoptera span a wide range of GC-content in non-coding regions (genome-wide average GCi ranging from 0.25 to 0.43), that strongly correlates with their average GC3 (from 0.32 to 0.71; Fig. 1B). This raises the question of how the tRNA pool evolved in these species in response to NSP changes. To investigate this point, we focused our analyses on the 26 Diptera and Lepidoptera species with a strong signal of translational selection.

In this dataset, we observed that the decoding of 11 NNA/NNG synonymous codon pairs (Glu, Gln, Lys, Val, Ala, Pro, Thr, Ser, both CTA/CTG and TTA/TTG pairs of Leu, and the AGA/AGG ‘duet’ of Arg) never involves wobble pairing: the two complementary isodecoder tRNAs (anticodons UNN and CNN, respectively) are systematically present altogether (Supplementary Fig. 8). Thus, for each of these 11 pairs, we were able to identify the ‘preferred’ isodecoder tRNA (*i.e.* the one with the highest gene copy number) in each species. We observed that the proportion of CNN anticodon among the 11 preferred tRNAs correlates positively with the average GCi (Fig. 6A). This implies that in species showing evidence of translational selection, tRNA gene copy number co-evolved in response to changes in NSP, consistent with the hypothesis that tRNA abundance is under selective pressure to match the demand in synonymous codon usage.

**Figure 6:**
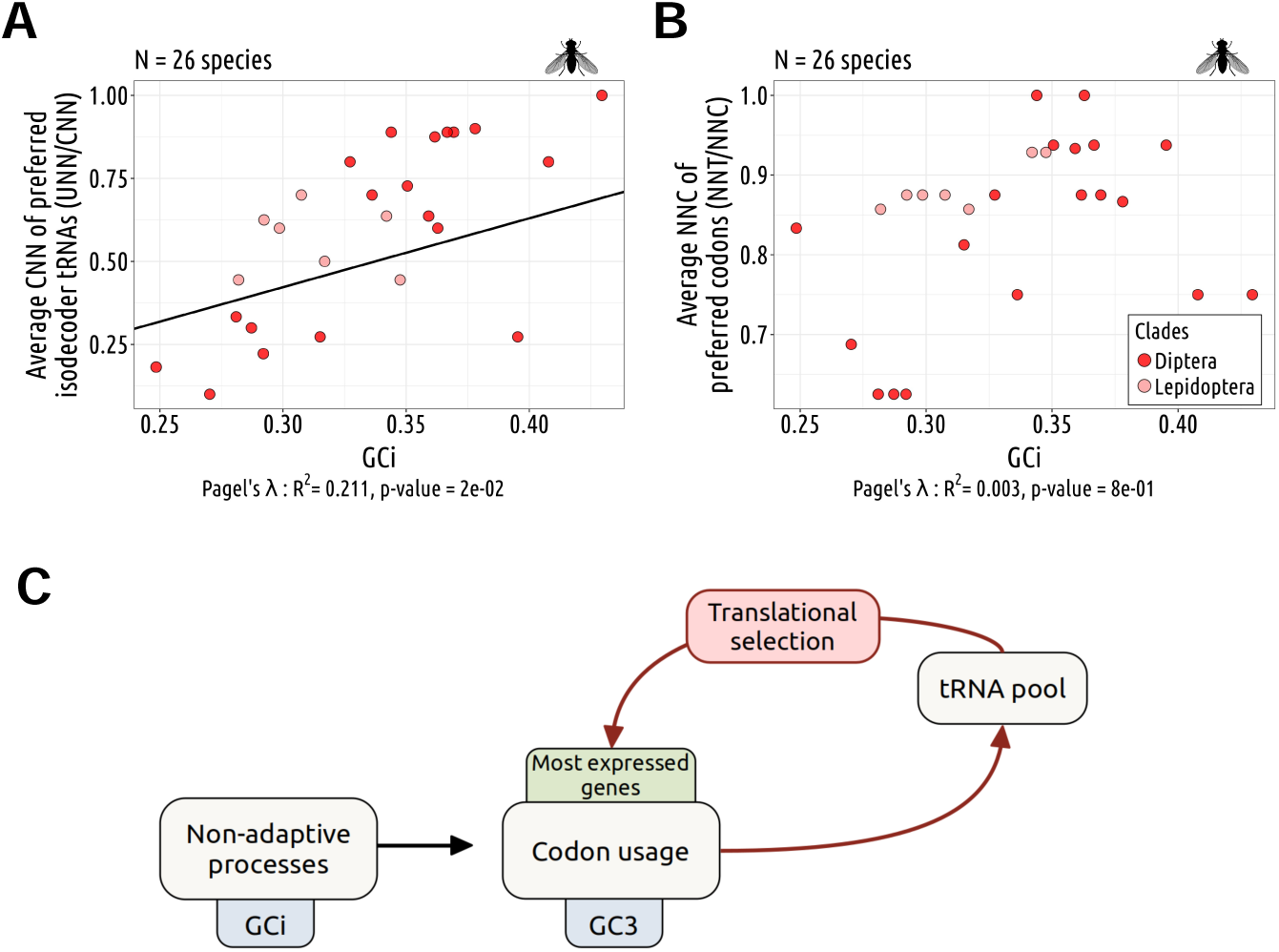
Genomic substitution patterns shape the tRNA pool. **A**: Relationship between the *per* species CNN fraction of preferred isodecoder tRNAs (corresponding to the most abundant tRNAs) among 11 NNA/NNG synonymous codon pairs and the gene average GC in introns (GCi), for Diptera (dark red) and Lepidoptera (light red). **B**: Relationship between the *per* species proportion of NNC preferred codons (the most overused codons in highly expressed genes compare to lowly expressed genes) among NNT/NNC synonymous codon pairs decoded by a single tRNA, along with the GCi. **A,B**: Pagel’s *lambda* model is used to take into account the phylogenetic structure of the data in a regression model (black line if significant). **C**: Hypothetical schemes explaining how synonymous codon usage can be shaped conjointly by translational selection and by neutral substitution patterns.

### Variation in optimality between wobble and Watson-Crick pairing

NNT/NNC synonymous codon pairs (N=16) are generally decoded by a single isodecoder tRNA (Fig. 3B, Supplementary Fig. 2). Among dipters and lepidopters, this is the case for 94% of the 416 NNT/NNC synonymous codons pairs analyzed (16 pairs *×* 26 species; Supplementary Fig. 8). In such cases, shifts in NSP cannot be compensated for by a change in the relative abundance of isodecoder tRNAs. Nevertheless, the affinity of tRNAs for their cognate codons can be changed by post-transcriptional modifications, and hence might evolve in response to the demand. To investigate whether such changes occur, it is necessary to identify which of the two codons is best decoded by this unique isodecoder tRNA. For this, we relied on the fact that in species that are subject to translational selection, codons that are more efficiently decoded show a higher prevalence in highly expressed genes compared to lowly expressed ones (these codons will hereafter be referred to as preferred codons). Two sets of NNT/NNC synonymous codons pairs can be distinguished: the 7 pairs corresponding to the amino acids with duet codons (Phe, Cys, Tyr, Asp, His, Asn and the AGT/AGC ‘duet’ of Ser), and the 9 pairs from amino acids with triplet (Ile) or quartet codons (Val, Gly, Ala, Pro, Thr, Leu, Arg, Ser). For NNT/NNC duets, when a single tRNA is present (95% of cases), it is always the GNN-tRNA, and in 99% of cases it is the NNC codon, decoded through Watson-Crick pairing that is preferred. For 8 of the 9 other pairs, when a single tRNA is present (94% of cases), it is always the ANN-tRNA, the only exception being Gly (GNN-tRNA). For Gly, the GGT codon, decoded via wobble pairing, is preferred to the GGC codon in 84% of species. For the other pairs (decoded by ANN-tRNA) there is more variability: the NNC codon (wobble pairing) is preferred in 79% of species, whereas the NNT codon (watson-crick pairing) is preferred in the others. These observations indicate that when a single tRNA is present for two codons, it is not systematically the one with watson-crick pairing that is the most efficiently translated. Furthermore, although the NNC codon tends to be preferred to the NNT codon (except for Gly), there are some variation across species, notably for those decoded by a ANN-tRNA (Supplementary Fig. 9). We computed in each species the proportion of NNC preferred codons among NNT/NNC synonymous codon pairs decoded by a single tRNA. Interestingly, the species showing the highest proportion of NNT preferred codons are the ones with the lowest genomic GC content (Fig. 6B). Thus, it appears that the relative affinity of ANN-tRNAs for the NNT or NNC codon can evolve in response to the demand.

These observations suggest a straightforward model to explain variation in the set of optimal synonymous codons across species (Fig. 6C). First, variation in mutational patterns or in the intensity of gBGC can lead to changes in the base composition of genomes, thereby directly shifting the codon usage of genes. Given that translational selection is a weak force, most genes are affected by this shift. This results in a change in the codon demand, and hence induces a selective pressure to change the pool of tRNA (both in terms of abundance and of affinity for their cognate codons). In turn, translational selection will modify the codon usage of highly expressed genes to match the new set of tRNAs, thereby reinforcing the selection on the tRNA pool to match the codon demand, and resulting into the coadaptation of tRNA-pool and codon usage.

### Weak translational selection in species with large intra-genomic variability in neutral substitution patterns

An implicit assumption of the above co-adaptation model is that all genes of a given genome are affected by similar neutral substitution patterns. There is evidence however that some genomes are subject to heterogeneous neutral substitution patterns. Notably, in mammals and birds, variation in recombination rates along chromosomes induces a strong heterogeneity in GC-content, driven by gBGC (Duret and Galtier, 2009). This process accounts for 70% of the variance in synonymous codon usage among human genes (Pouyet *et al*., 2017). Thus, in these species, the synonymous codon usage of a given gene essentially depends on the base composition of the genomic region where it resides, as shown by the strong correlation observed between GC3 and GCi across human genes (Fig. 1C). It is important to notice that these regional variations in GC-content affect all genes, even those that are widely expressed. To illustrate this point, we analyzed the codon usage of 2,249 human housekeeping genes (defined as genes that are in the top 20% most highly expressed genes in at least 75% of the tissues). The distribution of GC3 in housekeeping genes shows a very strong heterogeneity (GC3 ranging from 25% to 95%; Fig. 7A), as strong as in the entire gene set (Fig. 7B). Housekeeping genes are involved in basal function and have to be expressed at high level in most cell types. This implies that in any given cell, there is a strong heterogeneity of the codon demand. Such a situation is predicted to hinder the co-adaptation between the tRNA pool and codon usage: any increase in the abundance of a given tRNA (say decoding a GC-ending codon) is expected to be beneficial for the translation of GC-rich genes, but detrimental for the translation of the GC-poor ones (and vice versa for a tRNA decoding an AU-ending codon). Hence, the selective pressure imposed by the heterogeneous codon demand is expected to maintain a balanced tRNA pool, able to decode GC-rich genes as well as GC-poor genes. In turn, the presence of a balanced tRNA pool should reduce the difference in translational efficiency between synonymous codons, and hence is expected to decrease the intensity of translational selection. Thus, genomes that are subject to heterogeneous neutral substitution patterns are expected to be less subject to translational selection.

**Figure 7:**
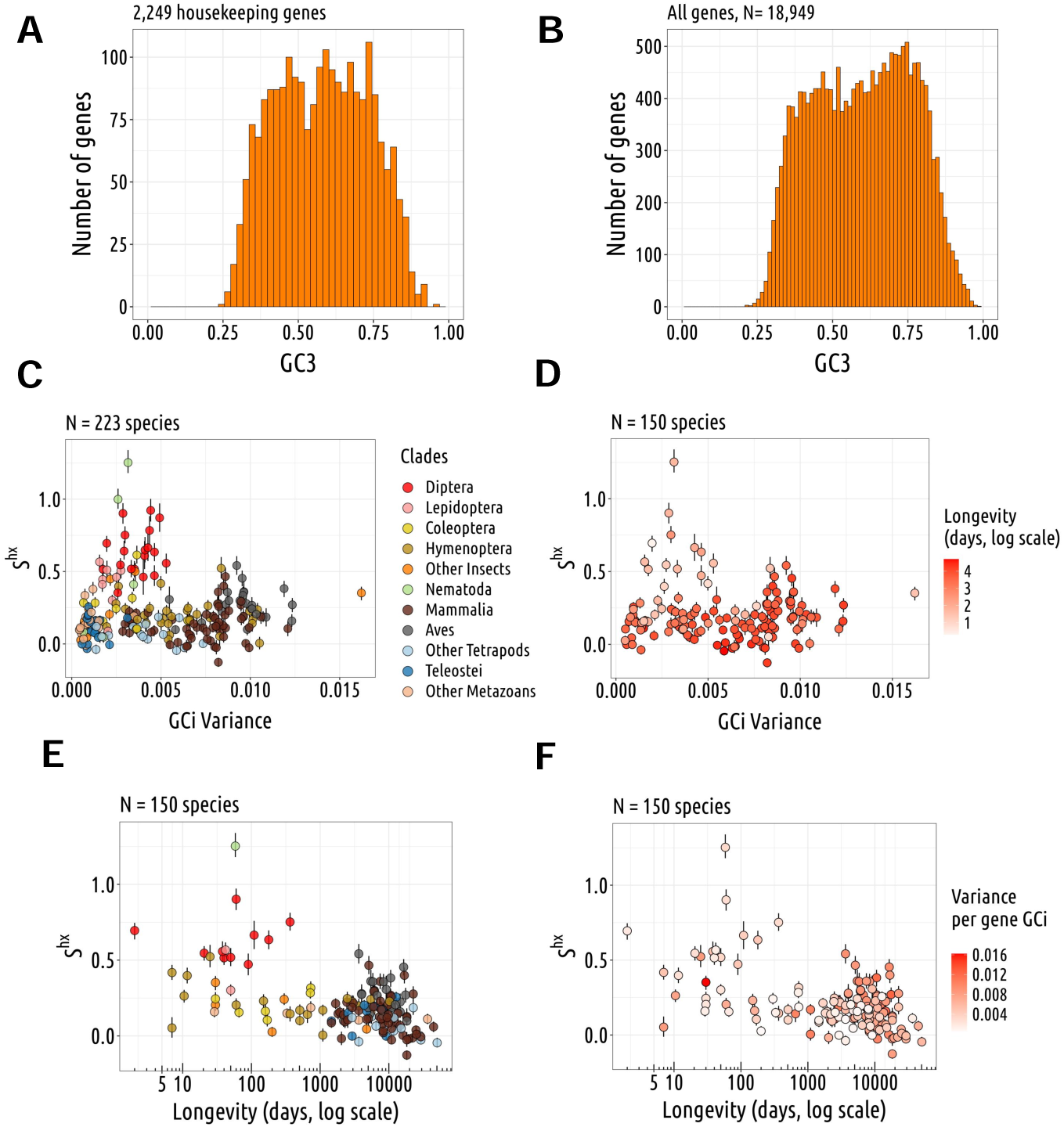
Impact of intra-genomic variability in neutral substitution patterns on translational selection. **A**: Distribution of GC3 across human housekeeping genes (identified based on gene expression data from 27 healthy tissues, extracted from (Pouyet *et al*., 2017). **B**: Distribution of GC content at the third position of codons (GC3) across all human genes. **C,D**: Relation between translational selection intensity *S*^hx^ and the gene GCi variance. **D**: Species are colored with a longevity gradient (log scale). **E,F**: Relation between translational selection intensity *S*^hx^ and longevity (days). **F**: Species are colored with a GC intron variance gradient (log scale) (N=223 species).

To test this prediction, we analyzed the relationship between the intensity of translational selection (*S*^hx^) and the intra-genomic heterogeneity in base composition (assessed by the variance in GCi across genes). We observed that all species with a strong signal of translational selection show a very small variance in GCi, while species with a high variance in GCi show relatively low *S*^hx^ (Fig. 7C). This is consistent with the hypothesis that intra-genomic heterogeneity in base composition precludes translational selection. However, species with a high variance in GCi mainly correspond to three clades (Mammal, Aves, Hymenoptera) that also have relatively small *N* _e_, and hence it is difficult to disentangle the impact of intra-genomic heterogeneity in base composition from that of drift (Fig. 7D). In any case, even though intra-genomic heterogeneity in base composition might explain the weakness of translational selection in some species, there must be some other factors that affect the intensity of translational selection. Indeed, among insect species predicted to have a *N* _e_ similar to that of dipters, many show a low *S*^hx^ despite a small variance in GCi (Fig. 7E,F).

## Discussion

### Impact of neutral substitution patterns on synonymous codon usage

Patterns of SCU vary widely across metazoan species, and are strongly correlated to the base composition of their non-coding regions (Fig. 1A). This implies that variation in codon usage across species are primarily shaped by differences in genome-wide neutral substitution patterns (driven by the underlying mutation pattern, by gBGC or both). NSPs vary not only across species, but also along chromosomes, and in some clades, such as tetrapods or hymenopters, this intra-genomic heterogeneity of NSPs is a major determinant of the variance in SCU among genes (Supplementary Fig. 1B). The fact that genome-wide patterns of SCU are strongly affected by NSPs does not exclude that it can also be shaped by a selective pressure favoring the use of translationally optimal codons. Indeed, in dipters and lepidopters, which both show clear evidence of translational selection, we observed that the tRNA pool evolves in response to changes in genome-wide NSPs (Fig. 6). Thus, variation in NSPs can lead to shifts in the translation apparatus, and thereby drive the evolution of SCU, not only in weakly expressed genes where codon usage is effectively neutral, but also in genes under strong translational selection. In other words, selective and non-adaptive models are not mutually exclusive, but it is important to take NSPs into account to be able to detect signatures of translational selection within genomes.

### Predicting translationally optimal codons

To quantify the intensity of translational selection in metazoans, we used a method based on standard population genetics equations, that infers the population-scaled selection coefficient (*S* = 4*N*_e_*s*) from the difference in optimal codon frequency between highly expressed genes and weakly expressed genes (Sharp *et al*., 2005; dos Reis and Wernisch, 2009). This method first requires to identify the set of optimal codons in each species. To predict optimal codons, previous studies generally searched for codons whose frequency increases with gene expression level (*e.g.* Duret and Mouchiroud (1999); dos Reis and Wernisch (2009)). One caveat, is that in some species, NSP varies with gene expression (Pouyet *et al*., 2017), which may therefore lead to errors in the inference of optimal codons. Furthermore, in that situation, the method would systematically overestimate *S* for codons that are favored by NSPs in highly expressed genes. This probably explains why their estimates of *S* in human and mice (respectively 0.51 and 0.22) are higher than ours (respectively 0.06 and 0.14). To limit this bias, we sought to predict optimal codons from the tRNA pool available in each species. For this, we estimated the abundance of each tRNA based on its gene copy number in the genome. In most species, we observed a strong correlation between the number of iso-acceptor tRNA gene copies and the frequency of their cognate amino acid in the proteome (Fig. 2B). These strong correlations are consistent with the fact that cellular tRNA abundance is highly constrained to match the amino acid demand, and indicate that tRNA gene copy number is a good proxy to infer tRNA abundance, in agreement with previous experimental evidence from a limited set of species (Behrens *et al*., 2021). Based on our estimates of the tRNA pool, we predicted two sets of putative-optimal codons (POCs): for amino acids for which more than one iso-decoder tRNA is available, optimal synonymous codons were defined as those decoded by the most abundant tRNA (POC1); for amino acids encoded by NNC/NNU duet codons and with one single iso-decoder tRNA (GNN), the NNC codons were predicted to be optimal (POC2), based on previous studies showing that the wobble pairing GNN:NNU was less efficient than the Watson-Crick pairing GNN:NNC (Stadler and Fire (2011); Chan *et al*. (2017); Fig. 3B).

Several lines of evidence indicate that our predictions of translationally optimal codons are accurate. First, our sets of POCs are consistent with previous predictions: among the optimal codons that had been inferred based on differential usage between highly and lowly expressed genes in *C. elegans* (N=26) and in *D. melanogaster* (N=25) (Duret and Mouchiroud, 1999), 88.4% and 88.0% respectively match with our POCs (25 POCs in *C. elegans* and 27 in *D. melanogaster*). Second, we observed that although the definition of POC1 and POC2 relies on very different principles, the two sets of codons show very similar signatures of translational selection (Fig. 4C). Furthermore, the analysis of substitution patterns and polymorphism in *Drosophila melanogaster* confirmed that selection favors POC alleles over non-POC alleles in highly expressed genes (Supplementary text 1). Finally, optimal codons are also expected to be selected for their impact on translation accuracy: consistent with this prediction, we observed that within genes, POCs are enriched at the more constrained amino-acid sites (Supplementary text 2).

### Variation in the intensity of selection in favor of translationally optimal codons across metazoans

For each species, we measured the frequency of optimal codons (combining POC1 and POC2) in highly expressed genes (top 2%), to estimate the population-scaled selection coefficient in favor of translationally optimal codons (*S* = 4*N*_e_*s*), using weakly expressed genes as a reference to account for the NSP (Sharp *et al*., 2005). Across the 223 species, the highest values of *S* are observed in *Caenorhabditis* nematodes (*S* = 1.25 in *C. elegans* and *S* = 1.00 in *C. nigoni*). We also found a clear signal of translational selection in diptera (mean *S* = 0.63, N=19 species), and to a lesser extent in lepidoptera (mean *S* = 0.41, N=8 species). Overall, estimates of *S* are weaker in other clades (Fig. 5). The weakness of translational selection in vertebrates (mean *S* = 0.15, N=119 species) was a priori expected given that these organisms tend to have relatively small *N* _e_. But what is surprising is that most of the invertebrates species show very weak translational selection. Besides *Caenorhabditis* and dipters, our dataset included 83 invertebrate species covering a wide range of clades (58 other insects, 12 other Ecdysozoa, 6 Spiralia, 4 Cnidaria, 3 Deuterostomia) that all show *S* values lower than that of dipters. This implies that the high values of *S* observed in *Caenorhabditis* and in dipters represent exceptions rather than the rule, and that translational selection is weak in most metazoan lineages.

If the selection coefficient in favor of optimal codons (*s*) was constant across metazoans, then *S* should scale linearly with *N* _e_. To test this prediction, we used silent-site polymorphism and germline mutation rate data (Lynch *et al*., 2023) to estimate the effective population size (*N_e_* = *π_s_*/4µ, hereafter noted 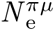) in 41 species. As expected *S* tends to increase with 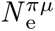, but the correlation is not significant after accounting for phylogenetic inertia (Fig. 5D). The weakness of the correlation might be due to the fact that these two parameters evolve on different time scales: 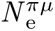 is indicative of the recent effective population size (on the order of *N* _e_ generations) and hence can change quite rapidly compared to *S*, that is estimated from the codon composition of genomes, resulting from a long-term accumulation of substitutions. This can explain why *C. nigoni* and *C. elegans* display similar values of *S*, despite a 75-fold difference in 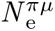 (respectively 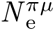 = 9.4 × 10^6^ and 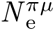 = 1.2 × 105). This difference in 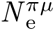 is due to the fact that *C. nigoni* is an outcrossing species, like most other *Caenorhabditis* species, while the *C. elegans* lineage evolved towards selfing hermaphroditism (Li *et al*., 2014; Vielle *et al*., 2016). This transition in reproductive mode is recent, and hence the SCU of *C. elegans* still retains the signature of strong translational selection inherited from its outcrossing ancestors. Thus, we can predict that the SCU of *C. elegans* is not at selection-mutation-drift equilibrium.

To further test the relationship between *S* and *N* _e_, we considered three additional parameters (longevity, body length and *dN/dS*), that are all correlated with 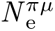 (Supplementary Fig. 7), but that are expected to reflect *N* _e_ over a longer time scale. A further interest of these proxies is that they can be estimated on much larger datasets (150 to 223 species). But here again we obtained similar results: *S* tends to increase with *N* _e_, but correlations are weak, marginally significant after accounting for phylogeny (Fig. 5A,B and C). The weakness of the correlation is mainly due to the fact that some species have a low *S*, despite life-history traits or *dN/dS* values indicative of a high *N* _e_.

Not only the correlations between *N* _e_ and *S* are weak, but also the range of variation in *S* appears to be quite limited compared to what would be expected given the variance in *N* _e_. For instance, the mean value of 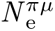 is about 15 times higher in diptera than in mammalia (based on respectively 6 and 41 species for which 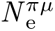 can be estimated; Lynch *et al*. (2023)). Yet, the mean value of *S* is only 5.3 times higher in diptera (mean *S* = 0.63, N=19 species) than in mammals (mean *S* = 0.12, N=65 species). Thus, the difference in *S* between diptera and mammals is smaller than what would be expected if *S* scaled linearly with *N* _e_.

One possible explanation for this discrepancy is that *S* is overestimated in mammals. As discussed by dos Reis and Wernisch (2009), the estimate of *S* is based on the assumption that NSPs are constant across genes, *i.e.* that the difference in optimal codon frequency between highly and weakly expressed genes is entirely due to translational selection. In reality, in some species, NSPs vary with gene expression (*e.g.* in humans Pouyet *et al*. (2017); Supplementary Fig. 10). To try to account for these variations, we measured the differences in POC frequency between highly and weakly expressed genes, controlling for differences in the corresponding triplet frequencies in introns. It is however possible that the base composition of introns is not a perfect predictor of NSPs, notably because introns are affected by indels and transposable elements, which are not allowed in coding regions. This is well illustrated by POC2 codons in humans, whose frequency clearly covaries with their non-coding controls, but with a wider amplitude in exons than in introns (Fig. 4A).

An alternative hypothesis is that the discrepancy might result from a strong heterogeneity in the fitness effect of synonymous mutations. Indeed, the analysis of synonymous polymorphism in *D. melanogaster* indicated that a majority of codons are under weak selection in favor of translationally optimal codon (*|N* _e_*s| ≈* 1), but that a small fraction (10%-20%) are under strong selection (*|N* _e_*s*| > 10; Machado *et al*. (2020). With a *N* _e_ value 15 times lower, the first class of codons is expected to evolve neutrally in mammals. But the second class of codons would still appear under effective translational selection, which might explain the small but non-null value of *S* measured in mammals.

Another unexpected observation is that many species predicted to have a high *N* _e_ (based on their LHTs or *dN/dS*) show very weak *S* (Fig. 5A,B and C). In some species, this could be explained by the heterogeneity of NSPs along chromosomes, inducing a strong variance in SCU that precludes a co-adaptation of the tRNA pool. This might be the case notably for some hymenopters, which, like tetrapods, are subject to gBGC (Wallberg *et al*., 2015) and present a very strong heterogeneity in NSPs (Fig. 7C). However, our dataset also includes some species with small *S* values, despite a high *N* _e_ proxy and homogenous NSPs. So finally, we are left with the conclusion that variation in *S* across metazoans are not driven simply by the drift barrier and by gBGC, but that they are also probably due to variation in s, the selection coefficient in favor of translationally optimal codons. There is evidence that in unicellular organisms, the selective force for the optimization of SCU is the maximization of cellular growth (Rocha, 2004; Sharp *et al*., 2005). It is possible that the selective pressure on cellular growth also vary across metazoans. Most *Caenorhabditis* species grow in ephemeral environments (rotting vegetation) and hence have been selected for their capacity to proliferate very rapidly (Cutter, 2015). Manthey *et al*. (2024) recently quantified growth rates in 33 insects. The only dipter present in their dataset (*Lucilia sericata*) is the species that presented the highest growth rate, 12 times higher than the average growth rate of other holometabole insects (N=10) and 52 times higher than the average growth rate of hemimetabole insects (N=22). If the *Lucilia sericata* is representative of other dipters, this might explain why translational selection is particularly strong in that clade compared to other insects. It is noteworthy that the two invertebrate species that have been historically used as model organisms (*D. melanogaster* and *C. elegans*) both belong to the very rare metazoan clades with clear evidence of translational selection. This might reflect the fact that they have been chosen as model organisms for the very reason that they can grow very fast in the lab.

## Materials & Methods

### Gene expression and data collection

The reference genome assemblies and genome annotations were downloaded from the National Center for Biotechnology Information (NCBI; Sayers *et al*. (2022). We obtained gene expression data for 257 metazoan species from GTDrift (Bénitìere *et al*., 2024), where gene expression levels (in Fragment *Per* Kilobase of exon *per* Million mapped reads, FPKM) were estimated over 8,362 RNA-seq samples. For each species we considered the *per*-gene median expression level across all analyzed RNA-seq samples. Additionally, a phylogenetic tree was retrieved from GTDrift to account for phylogenetic inertia (Bénitìere *et al*., 2024).

### tRNA gene annotation

The genomic coordinates of tRNA genes were extracted from the NCBI annotation file (N=44 species). For species in which tRNA annotations were not available (N=213 species), we predicted tRNA gene copies using the program tRNAscan-SE 2.0.12 (Nov 2022), with the -E option specifically designed for eukaryotic tRNA identification (Chan *et al*., 2021). To avoid tRNA pseudogenes, we retained only tRNA gene predictions with a score exceeding 55, threshold based on Chan *et al*. (2021).

### Analysis of codon usage

We analyzed codon usage only in genes detected as being expressed (>0 FPKM), so that to avoid erroneously annotated genes. For each gene, we counted codons of the longest annotated coding sequences (CDS). To control for variation in neutral substitution patterns, we measured the frequency of corresponding nucleotide triplets within intron (excluding the acceptor and donor splice sites that are highly constrained).

### Estimating Δ*POC^exp^*

We calculated Δ*POC^exp^* by subtracting the frequency of POCs in the 50% least expressed genes from the frequency of POCs in the top 2% most expressed genes. We then adjusted for this estimate by removing the observed differences in corresponding triplet frequencies within introns between the 2% most and the 50% least expressed genes.

### Estimates of effective population sizes

We retrieved proxies for the effective population size from the GTDrift data resource (Bénitìere *et al*., 2024), which included life history traits such as body length, longevity, and the ratio of non-synonymous to synonymous substitutions rate (*dN/dS*). It is expected that the genome-wide *dN/dS* ratio increases during prolonged periods of small *N* _e_, attributed to the fixation of slightly deleterious mutations (Ohta, 1992; Galtier, 2016).

GTDrift also provides measures of *N* _e_ based on the germline mutation rate (*µ*) and the level of neutral diversity (*π_s_*) published by (Lynch *et al*., 2023) and colleagues (*N_e_* = *π_s_*/4*µ*). We retrieved this estimate for 45 species, comprising 27 species within our dataset and 18 for which species from the same genus were available.

## Acknowledgements

Computational analyses were performed using the computing facilities of the CC LBBE/PRABI and the Core Cluster of the Institut Fraņcais de Bioinformatique (IFB) (ANR-11-INBS-0013). Silhouette images of taxonomic Families originate from PhyloPic developed and maintained by Mike Keesey available at https://www.phylopic.org/.

## Author contributions statement

LD, TL and FB designed the project. FB conceived the pipeline and conducted the analyses. LD, FB and TL wrote the manuscript.

## Funding

This work was funded by the French National Research Agency (ANR-20-CE02-0008-01 “NeGA”).

## Competing interests

The authors declare no conflicts of interest.

## Data and code availability

All processed data that we generated and used in this study, as well as the scripts that we used to analyze the data and to generate the figures, are available on Zenodo DOI: https://doi.org/10.5281/zenodo.12669922.

## Supplementary text

### 1 Selection favors optimal codons in highly expressed genes of *Drosophila melanogaster*

To further assess whether POCs are under selection in Diptera, we investigated patterns of polymorphism and substitution in *Drosophila*, based on a multiple genome alignment of three closely related species (*D. melanogaster, D. simulans, D. erecta*) and on single nucleotide polymorphism (SNP) data from 205 *D. melanogaster* individuals. We inferred the ancestral and derived state at each substitution or SNP, to distinguish synonymous changes corresponding to POC to non-POC mutations (PO*>nPO*) vs non-POC to POC mutations (nPO*>PO*) (see Materials & Methods; Supplementary Text Fig. 1A). To control for possible variation in local mutational patterns, we conducted a parallel analysis on triplets in intronic regions.

We observed that the rate of nPO*>PO* changes (number of nPO*>PO* changes/number of non-POC codons) increases with increasing gene expression level, while the rate of PO*>nPO* changes (number of PO*>nPO* changes/number of POC codons) decreased, both for SNPs and for substitutions(Supplementary Text Fig. 1B,D). Importantly, this trends is specific to coding regions, and is not observed for the corresponding triplets in introns (Supplementary Text Fig. 1C,E). These observations are consistent with the hypothesis that selection favors mutations leading to the incorporation of translationnally optimal codons in genes with high expression level.

### 2 Highly constrained amino acids are enriched in optimals codons

Synonymous codon usage is expected to be under selection not only for its impact on the speed of translation, but also on the accuracy of translation. For both traits, selective constraints are expected to vary among genes according to their expression level. One specific feature of selection for translation accuracy is that the strength of selection is also expected to vary among sites within a protein: selection on translation accuracy should be stronger at sites that are essential for the structure or function of the protein. To test this prediction, we analyzed within-gene variation in POC usage according to the level of constraint on amino acid sites. For this, we focused on a set of 976 orthologous genes, present in single copy in most metazoan genomes (BUSCO genes; Waterhouse *et al*. (2018)). For each protein of a given species, we classified its sites into four groups of equal size, according to their level of conservation across 293 metazoans (see Materials & Methods), and then measured the shift in POC frequency between its 25% most conserved sites and its 25% least conserved sites. Finally, we computed the average of these shift values over all proteins of this species (noted Δ*POC^cons^*). Given that the shift is computed within each gene, Δ*POC^cons^* measures variation in codon usage across sites that inherently have the same expression level.

**Supplementary Text 1:**
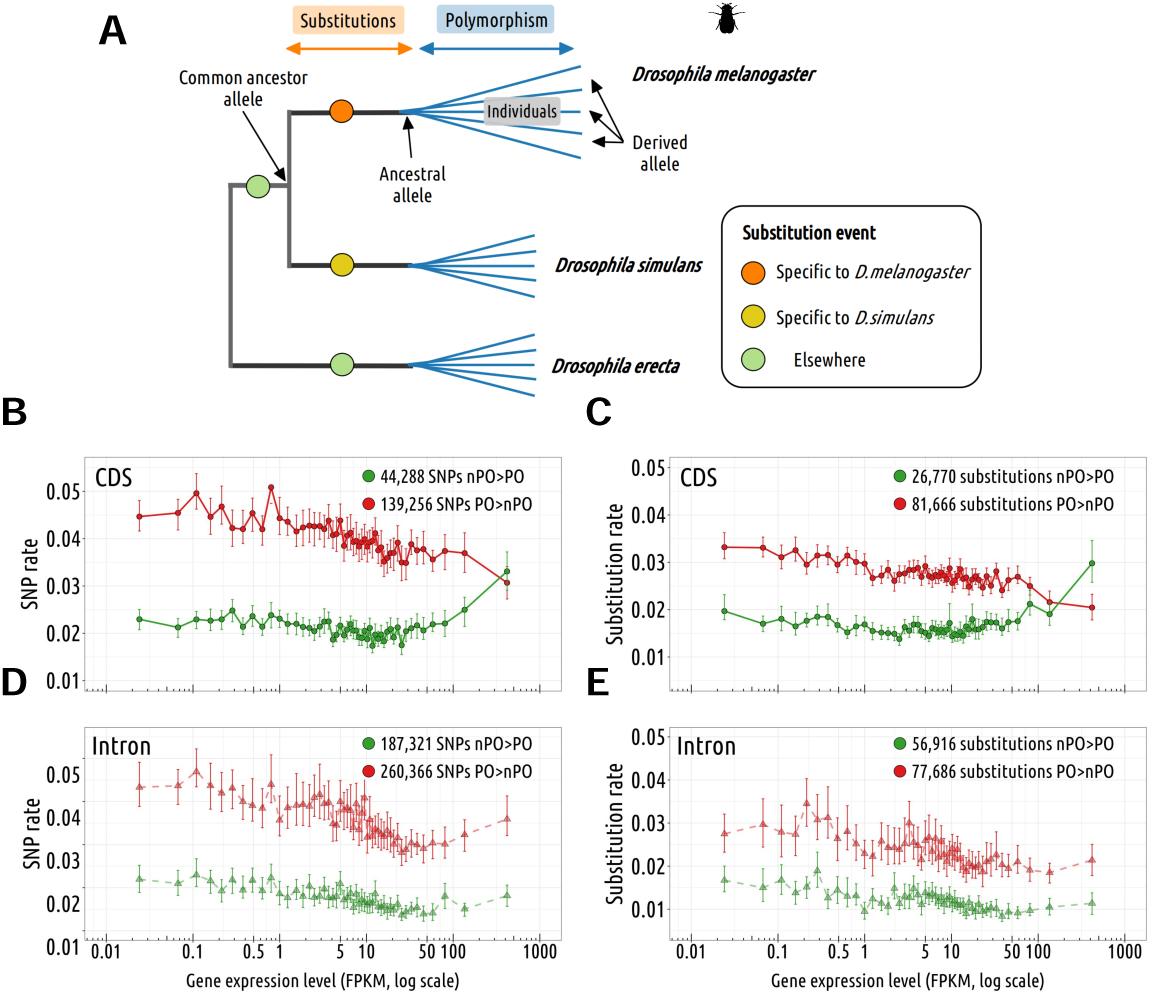
Selective pressure on non-POCs to POCs mutations. **A**: Schematic representation of the method used to identify SNPs and substitutions in *Drosophila melanogaster*. **B,C**: Rate variations of SNPs non-POC towards POC (green) and POC towards non-POC (red) with gene expression in CDS (**B**) and in intronic control (**C**). **D,E**: Rate variations of substitutions non-POC towards POC (green) and POC towards non-POC (red) with gene expression in CDS (**D**) and in intronic control (**E**). Error bars represent the 2.5th and 97.5th percentiles of values obtained by bootstrapping.

**Supplementary Text 2:**
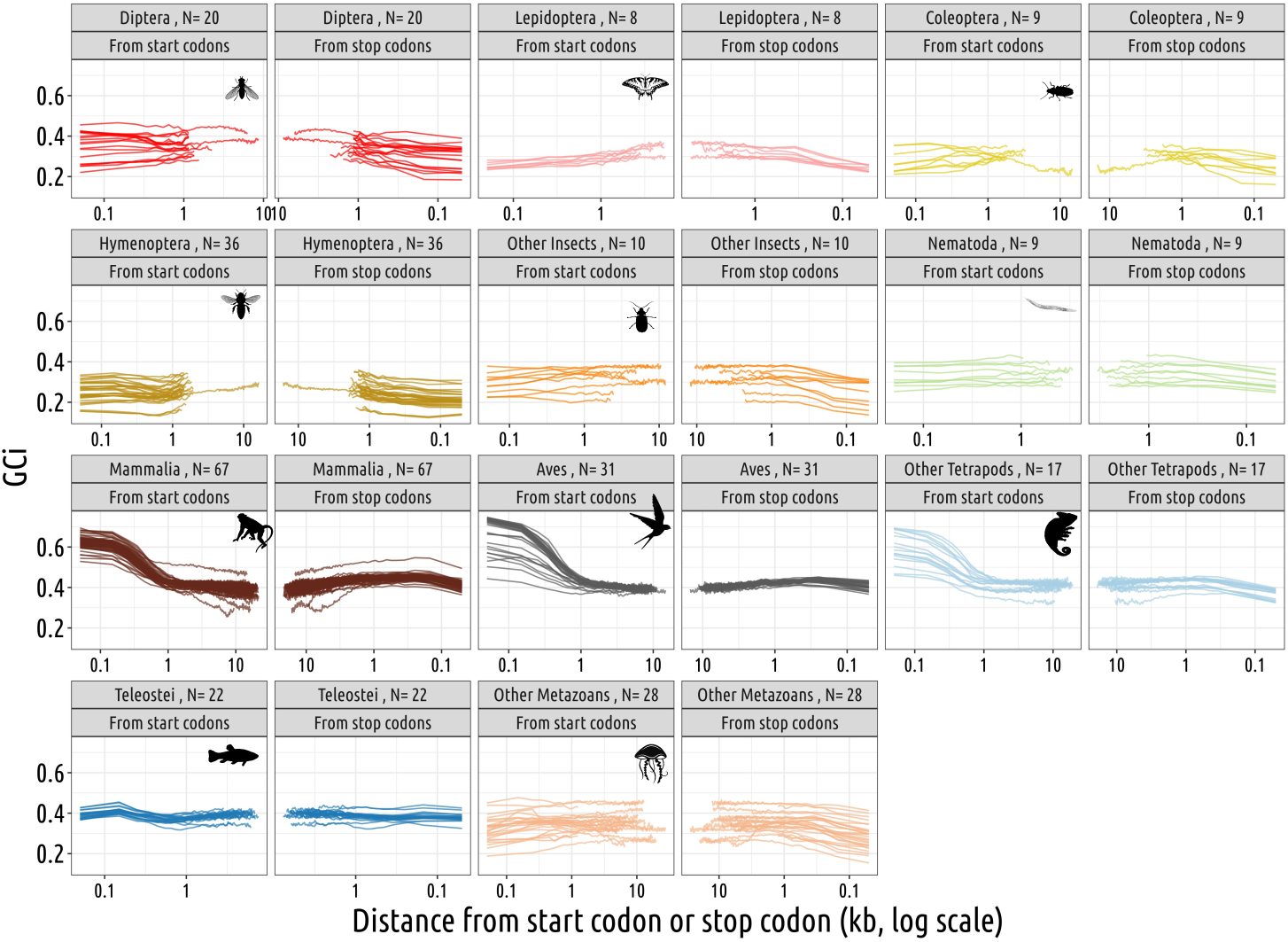
Non homogenous GC composition along genes for 11 clades. Measured of the GC composition in introns using 100 bp windows from the start codon and the stop codon (kb). Each clade of the study is represented by color, one line represents one species.

One difficulty however is that in vertebrates, many genes contain a GC-rich CpG island at their 5’ end (Deaton and Bird, 2011). The presence of a CpG island affects the base composition of the beginning of genes, up to about 1 kb, as illustrated by the analysis of the intronic GC content (Supplementary Text Fig. 2). As a result, the codon usage of the first exon is different than the rest of the coding region. Given that the N-termini of proteins evolve faster than their center (Bricout *et al*., 2023), this causes a spurious association between codon usage and variation in amino acid constraints along proteins. To avoid this bias, we measured Δ*POC^cons^* in vertebrates only on codons located beyond 1 kb of the start codon (in genomic coordinates). In other clades, the base composition of introns shows little variation along genes (Supplementary Text Fig. 2), and hence Δ*POC^cons^* was measured on the entire coding region.

In the *C. elegans*, the frequency of POC increased significantly between the least constrained and most constrained sites within proteins (from 48.5% on average to average 51.2%), whereas no variation was observed in humans (Supplementary Text Fig. 3A,B). Overall, Δ*POC^cons^* is positive in 73% of the tested species (refer to Supplementary Text Fig. 3C). As for Δ*POC^exp^s*, Δ*POC^cons^* shows substantial variation across clades, and is maximal for Diptera.

**Supplementary Text 3:**
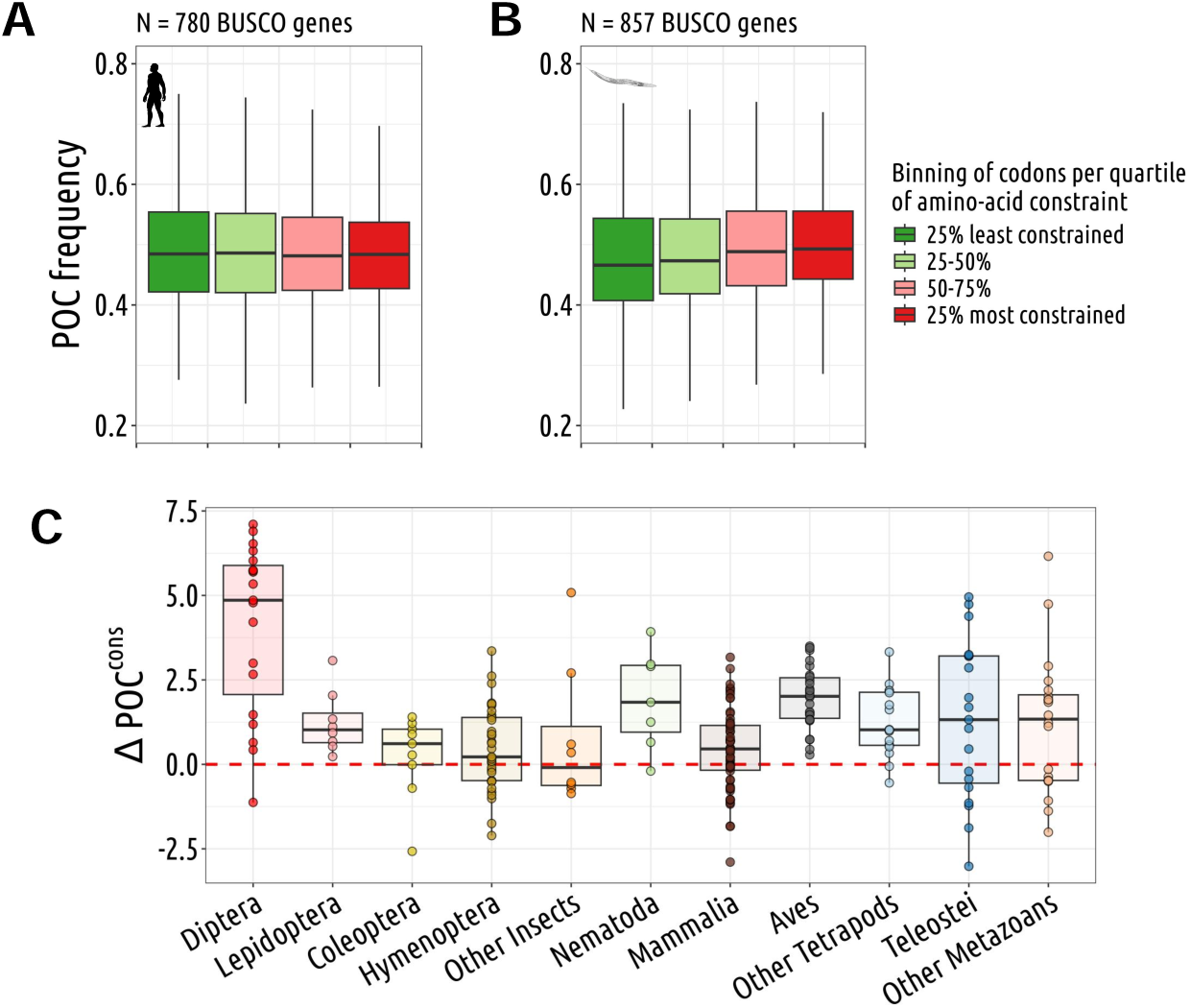
Highly constrained amino acids are enriched in POCs. **A,B**: We analyzed intragenic variation in codon usage within the set of BUSCO genes (N=976 genes). Within each gene, codons were divided into four groups of equal size, according to the level of constraint on the corresponding site in the protein. The level of constraint on amino-acid sites was estimated based on the proportion of gaps in the protein alignments across the 293 species. The frequency of POCs was calculated for each gene within each constraint group (from green to red boxplots). **A** represents *Homo sapiens*, and **B** represents *Caenorabditis elegans*. **C**: For each species (N=223 species), we measured the difference in POC frequency between the most constrained and least constrained groups. The plot show the distributions of this difference across different clades.

## Materials & Methods

### Variation in codon usage according to the level of constraints on amino acid sites

Multiple gene alignments of 976 BUSCO genes and 293 species were collected from the metazoa dataset alignment of GTDrift repository (Bénitìere *et al*., 2024). The level of constraint on amino acid residues was measured as the proportion of gaps at the corresponding site in the protein alignment. For each gene of each species, codons were binned into four groups of equal size, based on their level of constraint. In vertebrates, we focused on sites located beyond 1,000 base pairs downstream from the start codon (in the genomic sequence).

### Analysis of synonymous polymorphism and divergence

We used polymorphism data from the *Drosophila* Genetic Reference Panel 2 (DGRP2) (Mackay *et al*., 2012; Huang *et al*., 2014), where polymorphic sites have been identified from comparisons across 205 inbred lines of *Drosophila melanogaster*, downloaded from http://dgrp2.gnets.ncsu.edu/data/website/dgrp2.vcf. We converted the single nucleotide polymorphism (SNP) coordinates from the dm3 genome assembly to the dm6 assembly, with the liftOver utility (Hinrichs *et al*., 2006) of the UCSC genome browser, using a whole genome alignment between the two assemblies downloaded from https://hgdownload.soe.ucsc.edu/goldenPath/ dm3/liftOver/dm3ToDm6.over.chain.gz. We kept in the study 3,738,302 biallelic SNPs for which more than 181 individuals have been genotyped.

We then identified two sister species *Drosophila simulans* (from https://hgdownload.soe.ucsc.edu/gbdb/ dm6/liftOver/dm6ToDroSim1.over.chain.gz) and *Drosophila erecta* (from https://hgdownload.soe. ucsc.edu/goldenPath/dm6/liftOver/dm6ToDroEre2.over.chain.gzz) that we aligned against *Drosophila melanogaster* genome using liftOver utility (Hinrichs *et al*., 2006). We removed from the analysis genes located in regions where the multiple alignment was of poor quality, and genes that were not detected as being expressed (to avoid erroneous gene annotations). In case of alternative splicing, we retained the annotated transcript that encodes the longest protein. We used the program est-sfs release 2.04 (Keightley and Jackson, 2018) to polarize SNPs, *i.e.* to identify the ancestral allele and the derived allele. The same approach was applied for intron regions by studying nucleotide triplets.

We also used the multiple genome alignment of *Drosophila simulans*, *Drosophila erecta* and *Drosophila melanogaster* to identify substitutions by parsimony (see Supplementary Text Fig. 1A). To focus on fixed differences, polymorphic sites were excluded.

## Supplementary figures

**Supplementary Figure 1:**
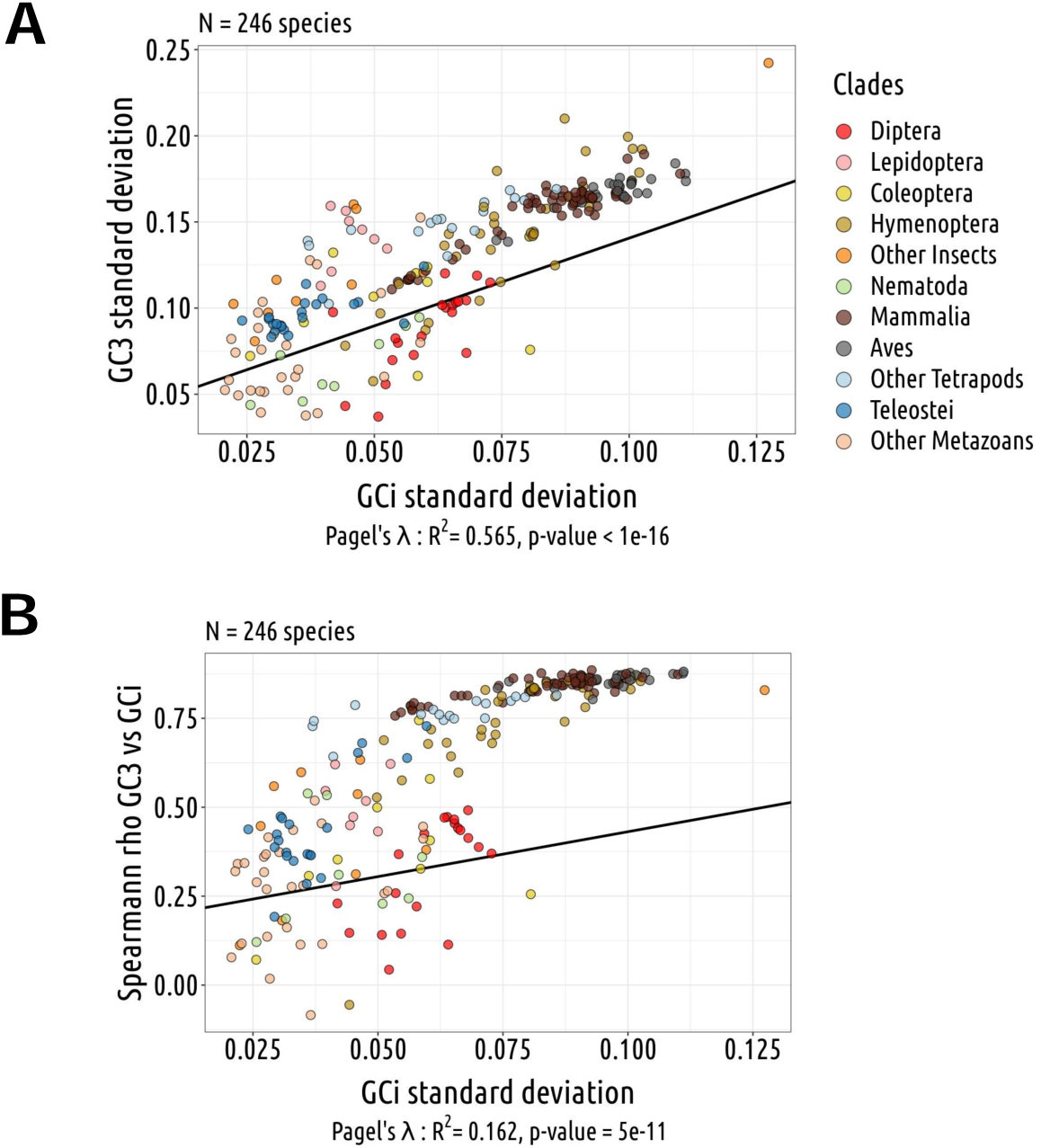
Intra-genomic variation in base composition. **F**or each species, we measured the GC-content of each gene, in introns (GCi) and at third codon positions (GC3). We then quantified the variation in GC3 and GCi among genes, and we measured the correlation between GC3 and GCi (Spearman correlation, rho). We then analyzed how these parameters covary across species. A: Relationship between the standard deviation of GC3 and of GCi. Species with a strong intra-genomic variability in GC-content in non-coding sequences (GCi) also display a strong heterogeneity of codon usage (GC3)**B**: Relationship between rho, and the standard deviation of GCi. In species with a strong variance in GCi, codon usage (GC3) is strongly correlated to the base composition of non-coding sequences (GCi).

**Supplementary Figure 2:**
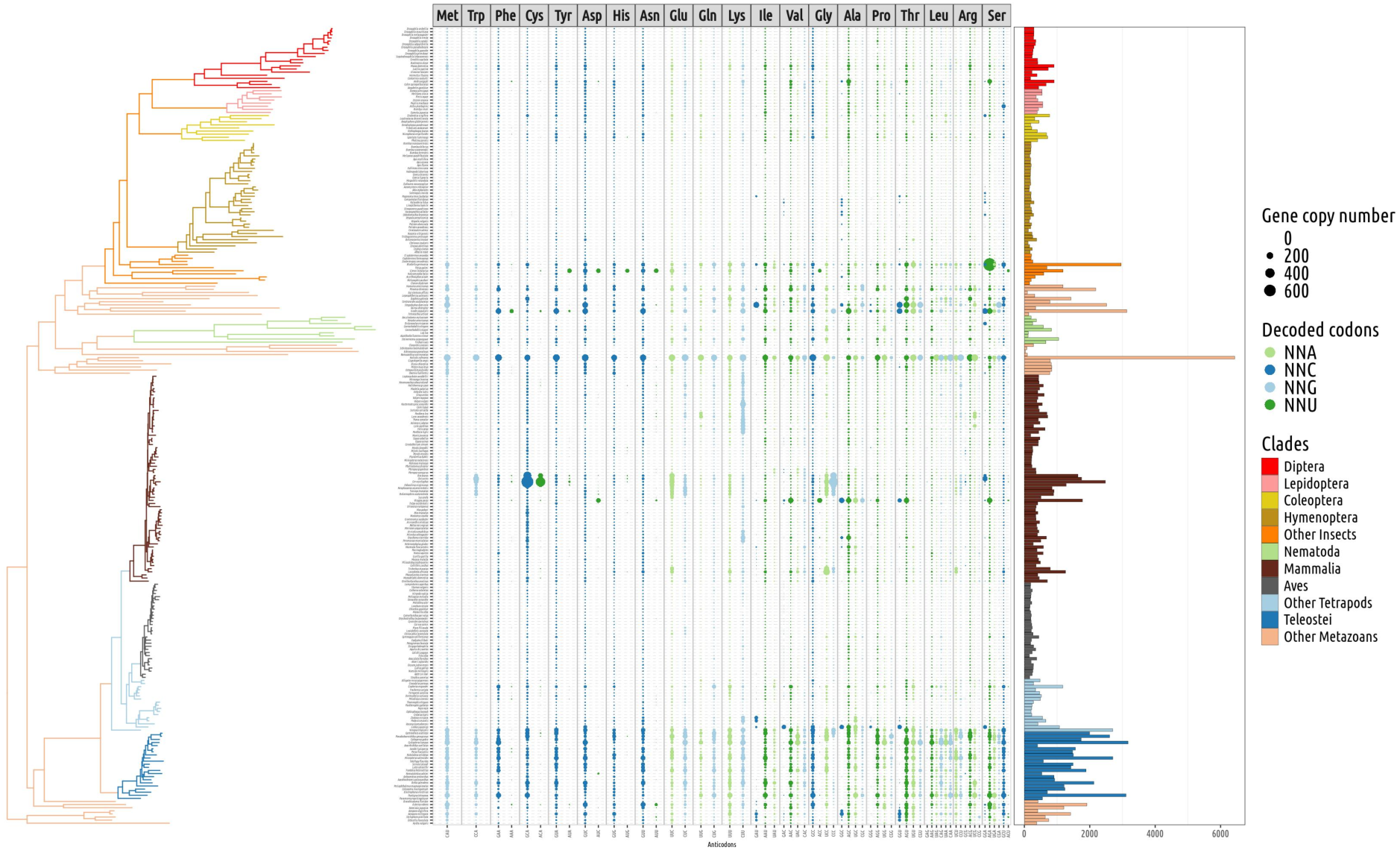
tRNA gene copy number. For the 257 studied species from left to right: phylogenetic tree, number of tRNA gene copies *per* anticodon (grouped *per* amino acid) and total number of tRNA gene copies.

**Supplementary Figure 3:**
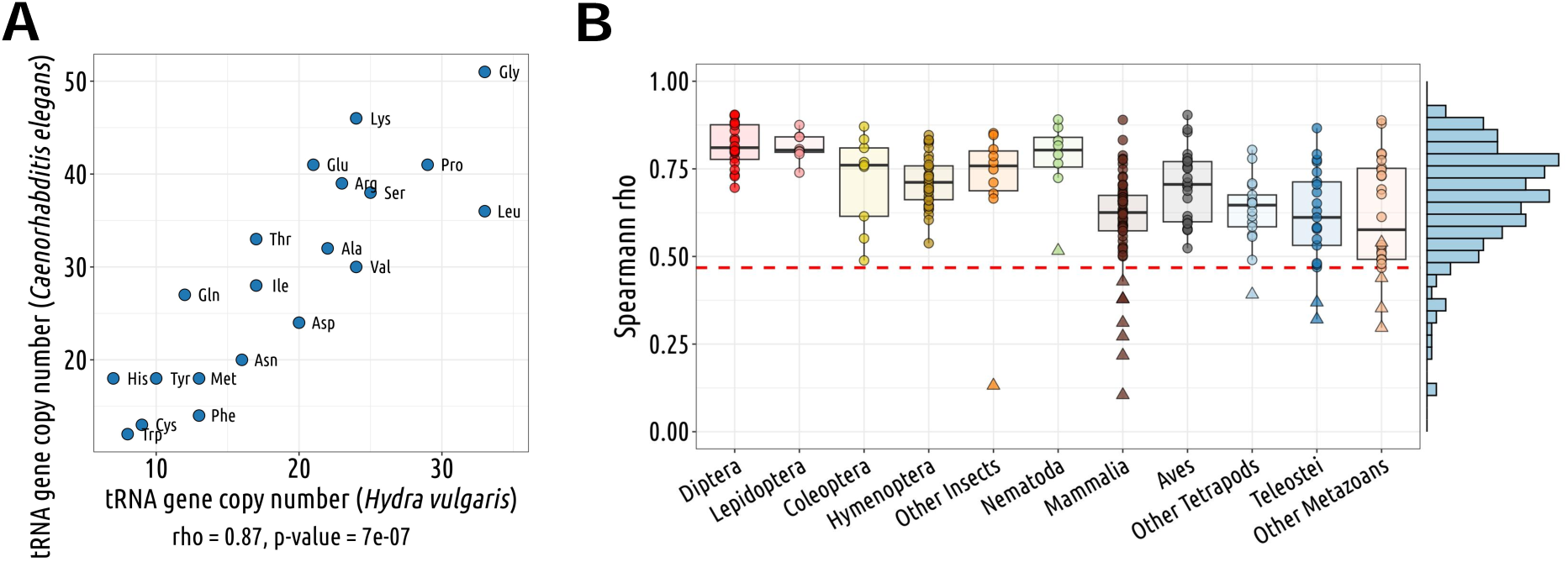
Conservation of the relative abundance of isoacceptor tRNA genes across metazoan genomes. **F**or each species, we counted the number of tRNA gene copies for each amino-acid, and measured the correlation with the corresponding counts in the genome of *Hydra vulgaris*, used as a reference. A: Representative example illustrating the conservation of the relative number of tRNA isoacceptor genes between *Caenorhabditis elegans* and *Hydra vulgaris*. Spearman’s correlation (rho) and the corresponding p-value are displayed under the graph. **B**: Distribution of this conservation index (rho between *Hydra vulgaris* and each focal species) across different clades (N=245 species). The red line denotes the threshold above which the p-value is less than 0.05.

**Supplementary Figure 4:**
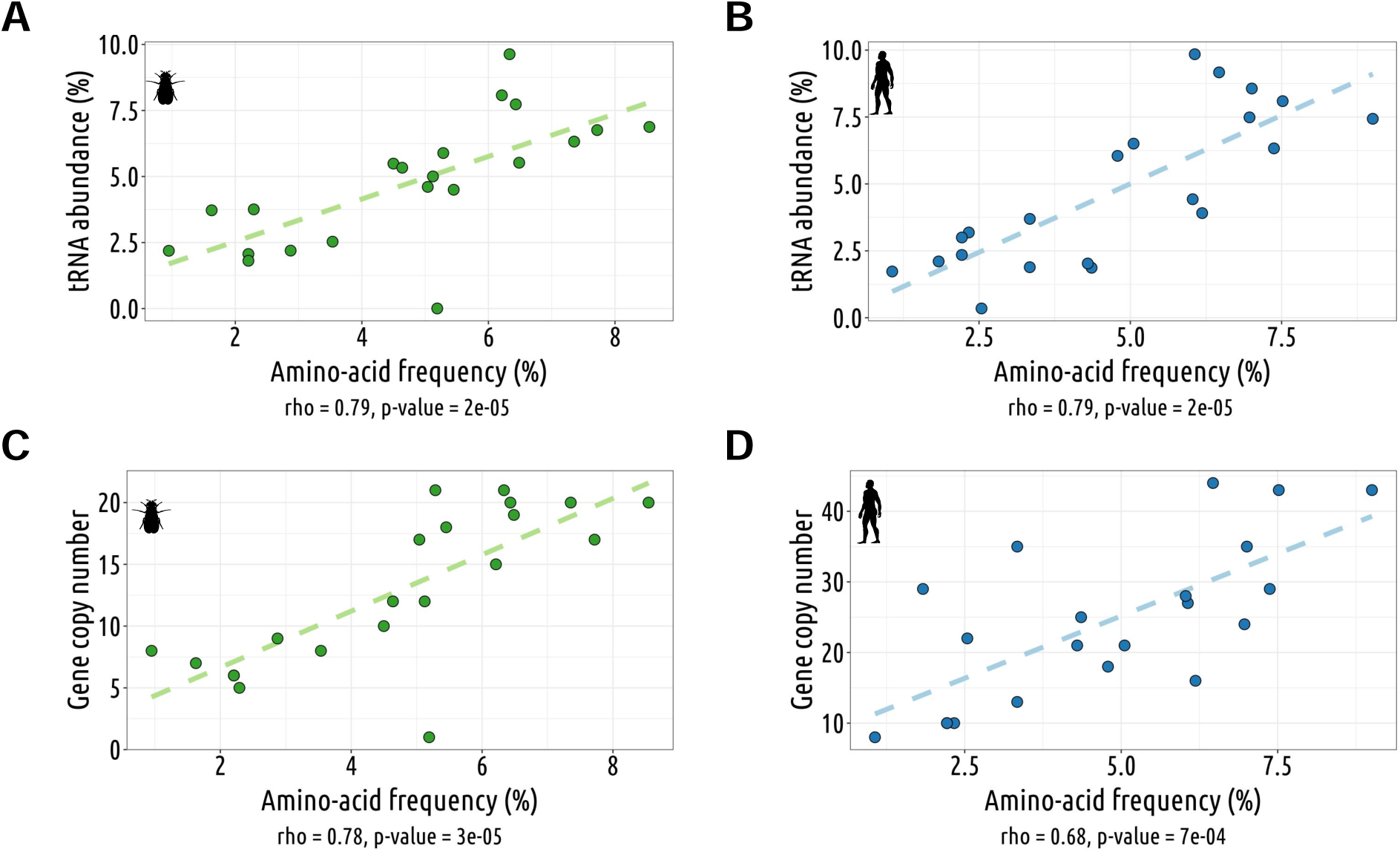
Relationship between the amino-acid demand and tRNA abundance: direct vs indirect estimates of tRNA abundance. The amino-acid demand corresponds to the frequency of amino acids in the proteome, weighted by gene expression levels. **A,B**: Relationship between amino-acid demand and direct measures of tRNA abundance from Behrens *et al*. (2021). **C,D**: Relationship between amino-acid demand and tRNA gene copy number (proxy for tRNA abundance). Left: *Drosophila melanogaster* ; Right: *Homo sapiens*

**Supplementary Figure 5:**
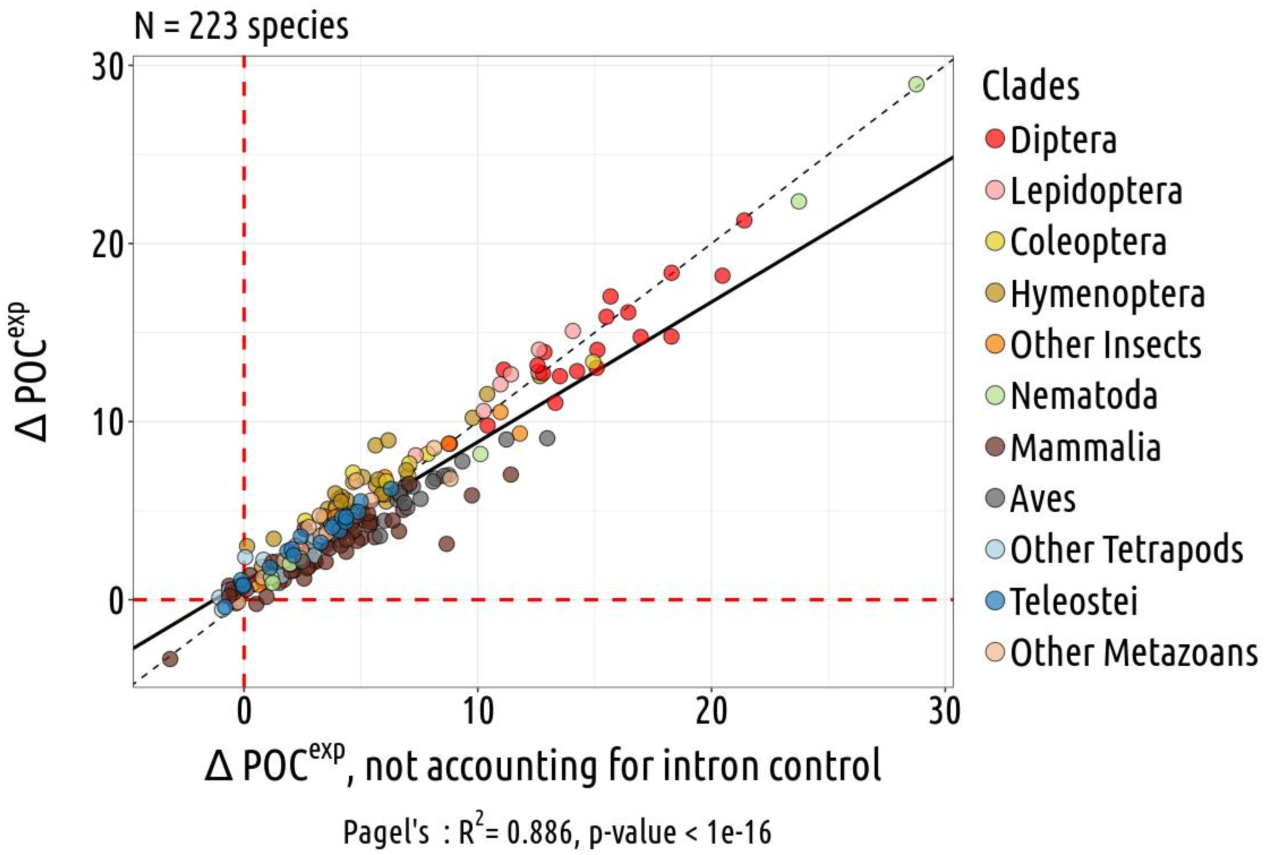
Quantifying the signal of translational selection, accounting or not for variation in intronic base composition. To investigate the impact of TS on synonymous codon usage, we measured the difference (noted Δ*POC^exp^*) between the frequency of POCs in the most expressed genes (top 2%), and the frequency of POCs in the 50% least expressed genes. In our main analyses, to account for intra-genomic variation in neutral substitution patterns, we computed Δ*POC^exp^*, controlling for variation in base composition within introns (see Materials & Methods). The comparison of Δ*POC^exp^* values measured with or without this control shows that this control step does not have a strong impact on our results. The black line represents the pagel’s *lambda* model, and the dotted line represents x=y.

**Supplementary Figure 6:**
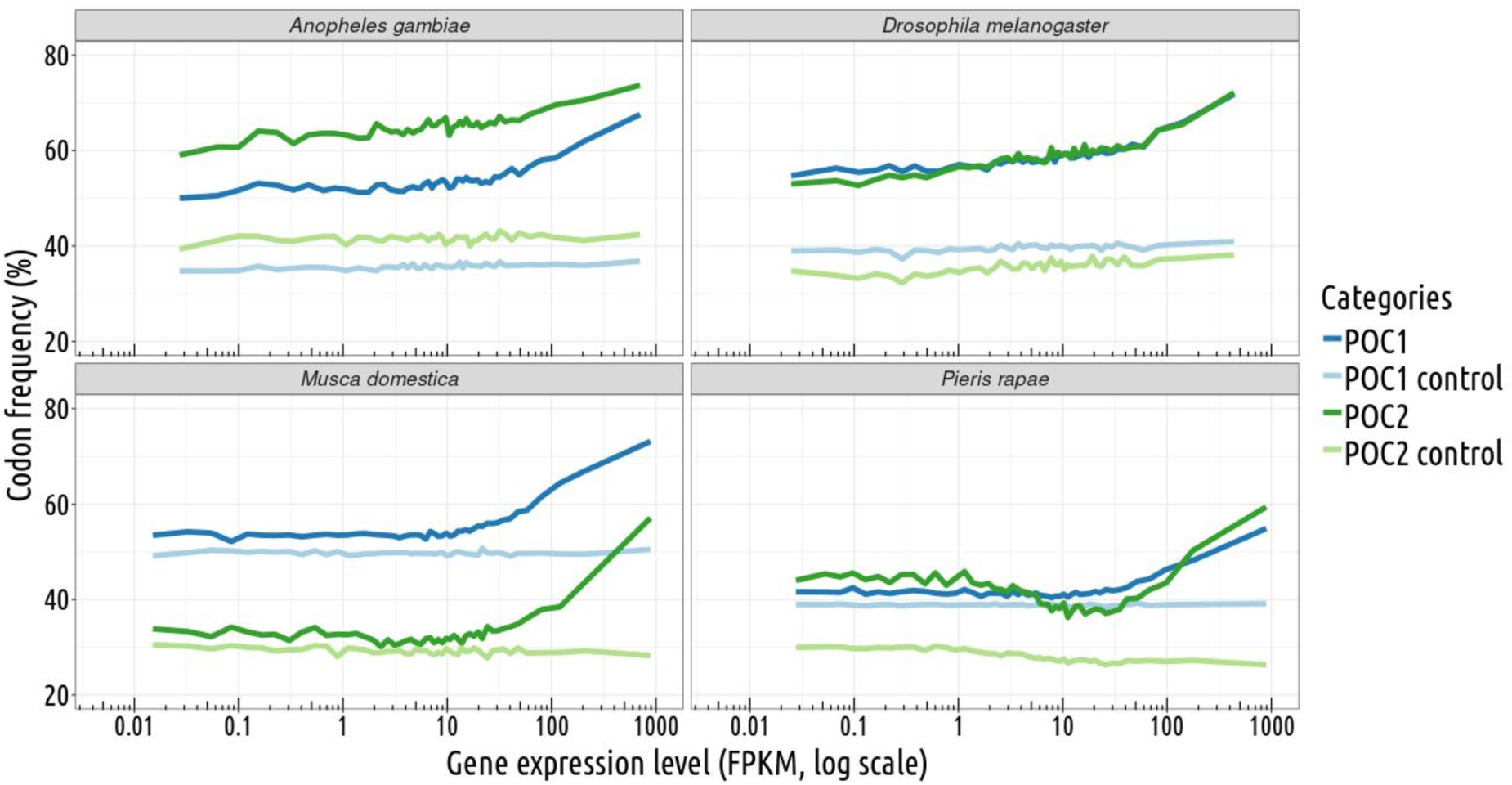
Relationship between the frequency of putative-optimal codons and gene expression level in 4 representative species. Variation in the proportion of POC within coding sequences (POC1: dark blue; POC2: dark green) according to gene expression level. To control for variations in neutral substitution patterns, we analyzed the frequency of corresponding triplets within introns (POC1 control: light blue; POC2 control: light green). Each point represents a 2% bin of genes.

**Supplementary Figure 7:**
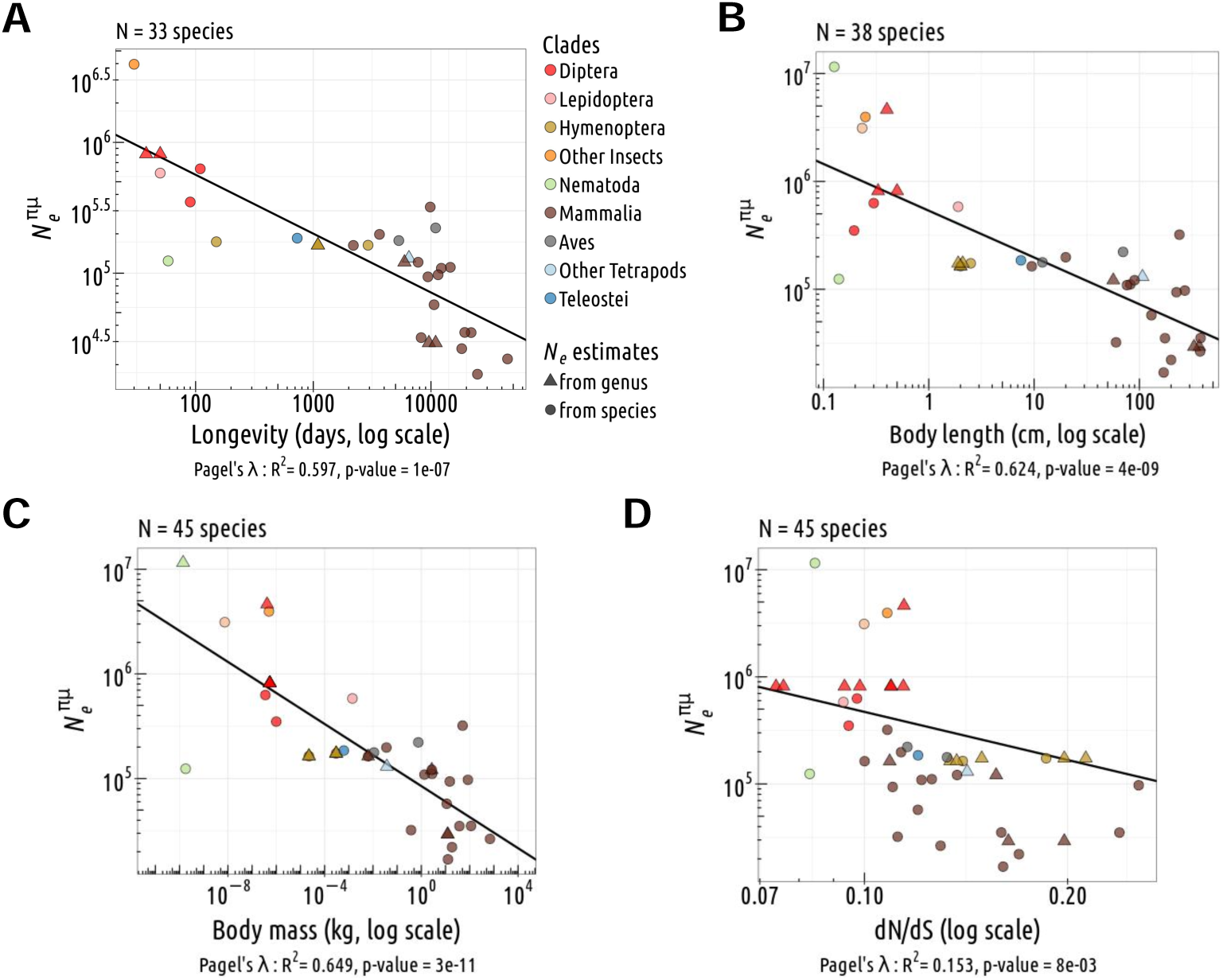
Relationship between different proxies of *N* _e_. Relationship between *N ^πµ^* (estimated based on germline mutation rate and on the level of silent polymorphism) and the longevity (**A**), body length (**B**), body mass (**C**), *dN/dS* (**D**). Pagel’s *lambda* model is used to take into account the phylogenetic structure of the data in a regression model (the regression line is displayed in black when the correlation is significant).

**Supplementary Figure 8:**
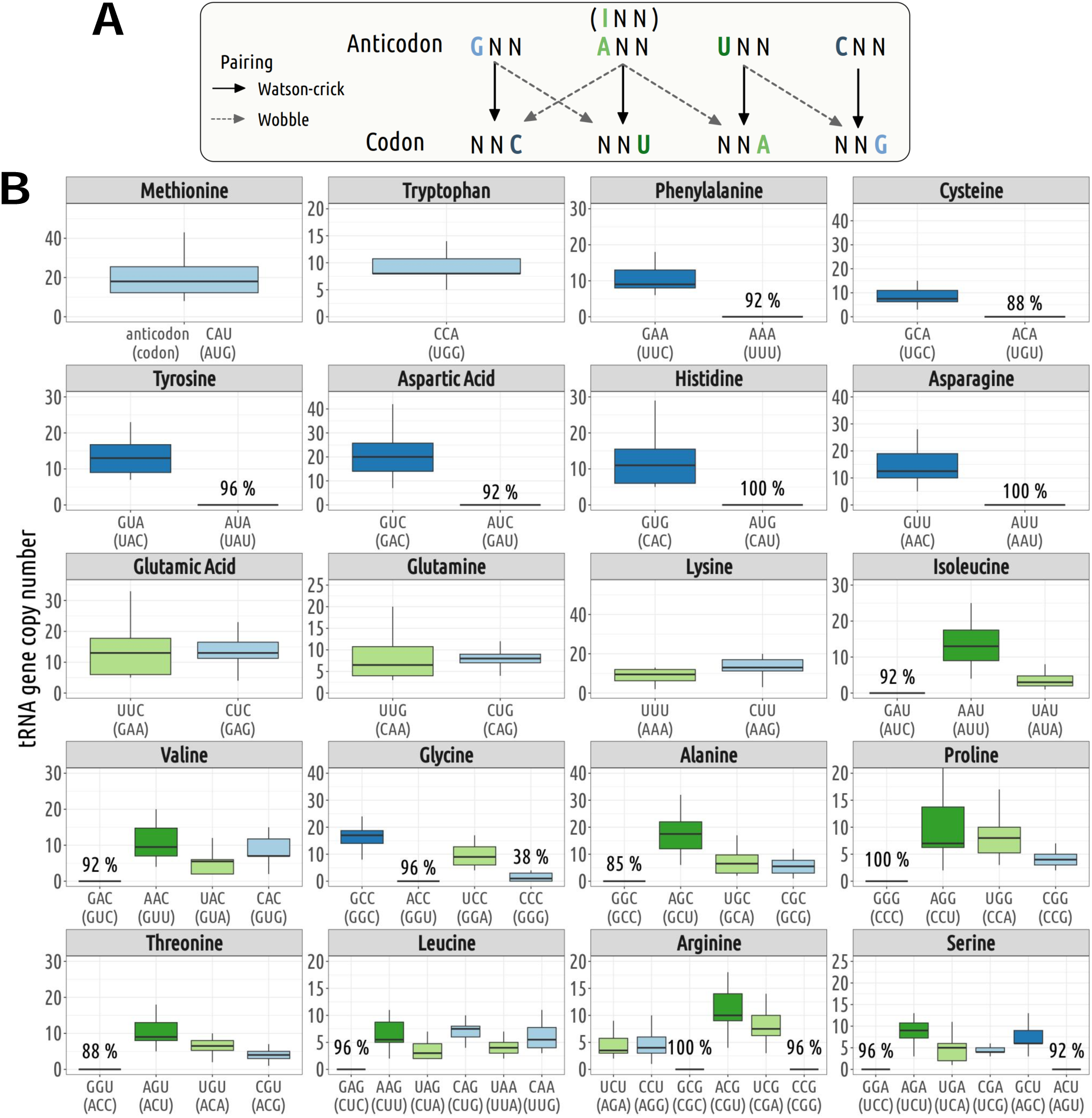
Codon-anticodon assignment for Lepidoptera and Diptera. In each genome, we used the rules proposed by Percudani (Percudani, 2001) to assign codons to their cognate tRNA: first, isodecoder tRNAs present in a given genome are assigned to their complementary (Watson-Crick) codons; then, the remaining codons (for which the genome does not contain any tRNA carrying the complementary anticodon), are inferred to be decoded by wobble pairing. **A**: Illustration of the various possible pairings: Watson-Crick and wobble pairing. **B**: Distribution of tRNA gene copy numbers across 26 species subject to translational selection (Lepidoptera and Diptera). The boxplot represents the median, interquartile range (box edges at 25th and 75th percentiles), and whiskers extending to the largest value no further than 1.5 times the interquartile range. For tRNA isodecoders that are absent from some genomes, the percentage of species lacking a tRNA gene copy is also indicated.

**Supplementary Figure 9:**
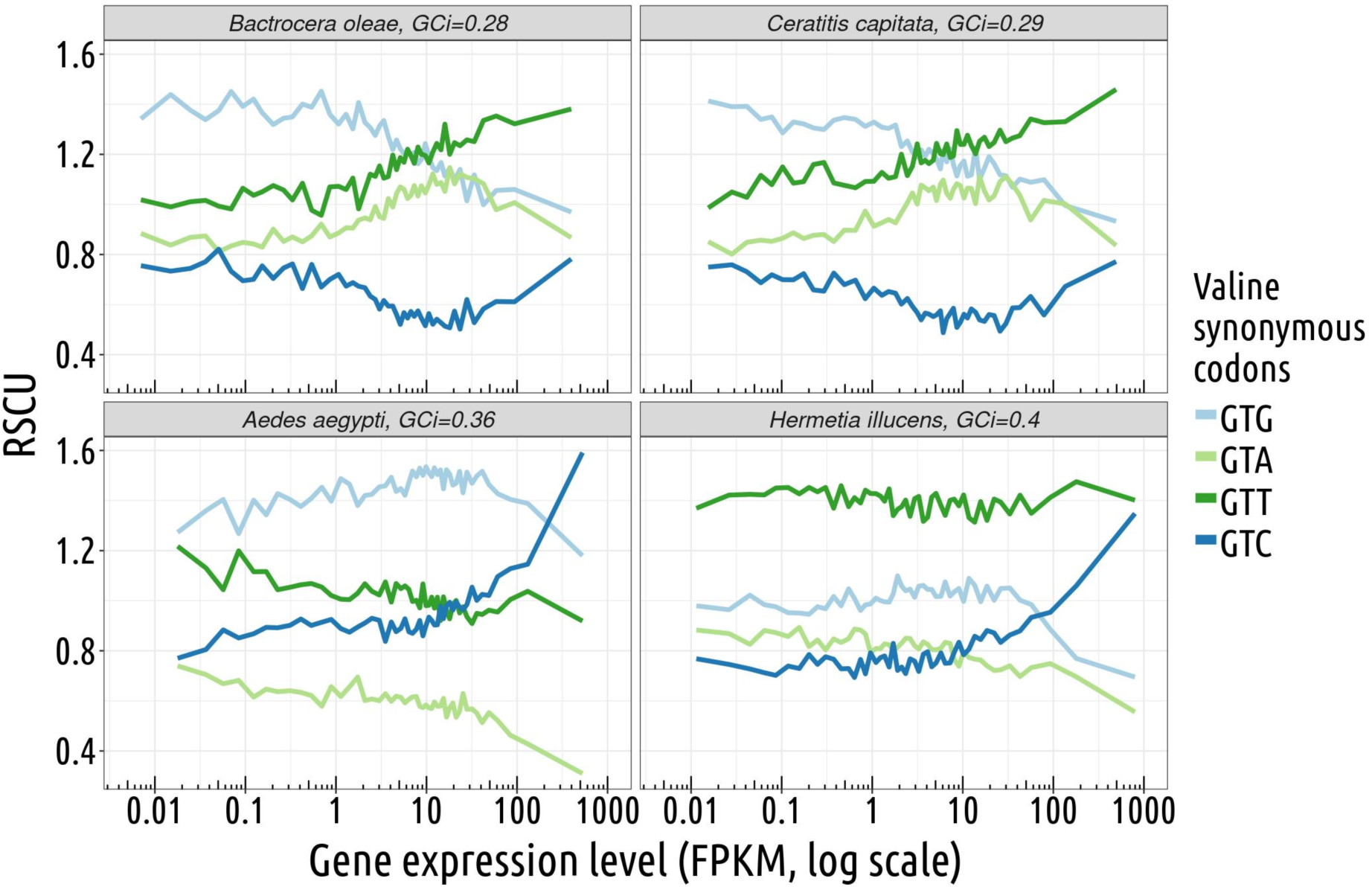
Variation in the optimality in Valine synonymous codons among four dipter species with different GC richness. The optimality of valine synonymous codons (GTG/GTA/GTT/GTC) can be assessed by analysing the relationship between their relative synonymous codon usage (RSCU) and gene expression level. In these four dipters, Valine has two POC codons (GTC and GTT) decoded by a single isodecoder tRNA (anticodon AAC). As expected, in all cases, the frequency of non-POC codons (GTG and GTA) decreases with increasing gene expression levels, which indicates that they are not optimal. In GC-poor genomes (*Bactrocera oleae*, *Ceratitis capitata*), GTT is used much more frequently than GTC, but both POCs increase in frequency with gene expression, which suggests that they both are optimal. In the two more GC-rich genomes (*Aedes aegypti* and *Hermetia illucens*), only the GTC codon shows an increase in frequency with expression, which suggests that GTT (decoded by Watson-Crick pairing) is less efficient than GTC (decoded by wobble pairing).

**Supplementary Figure 10:**
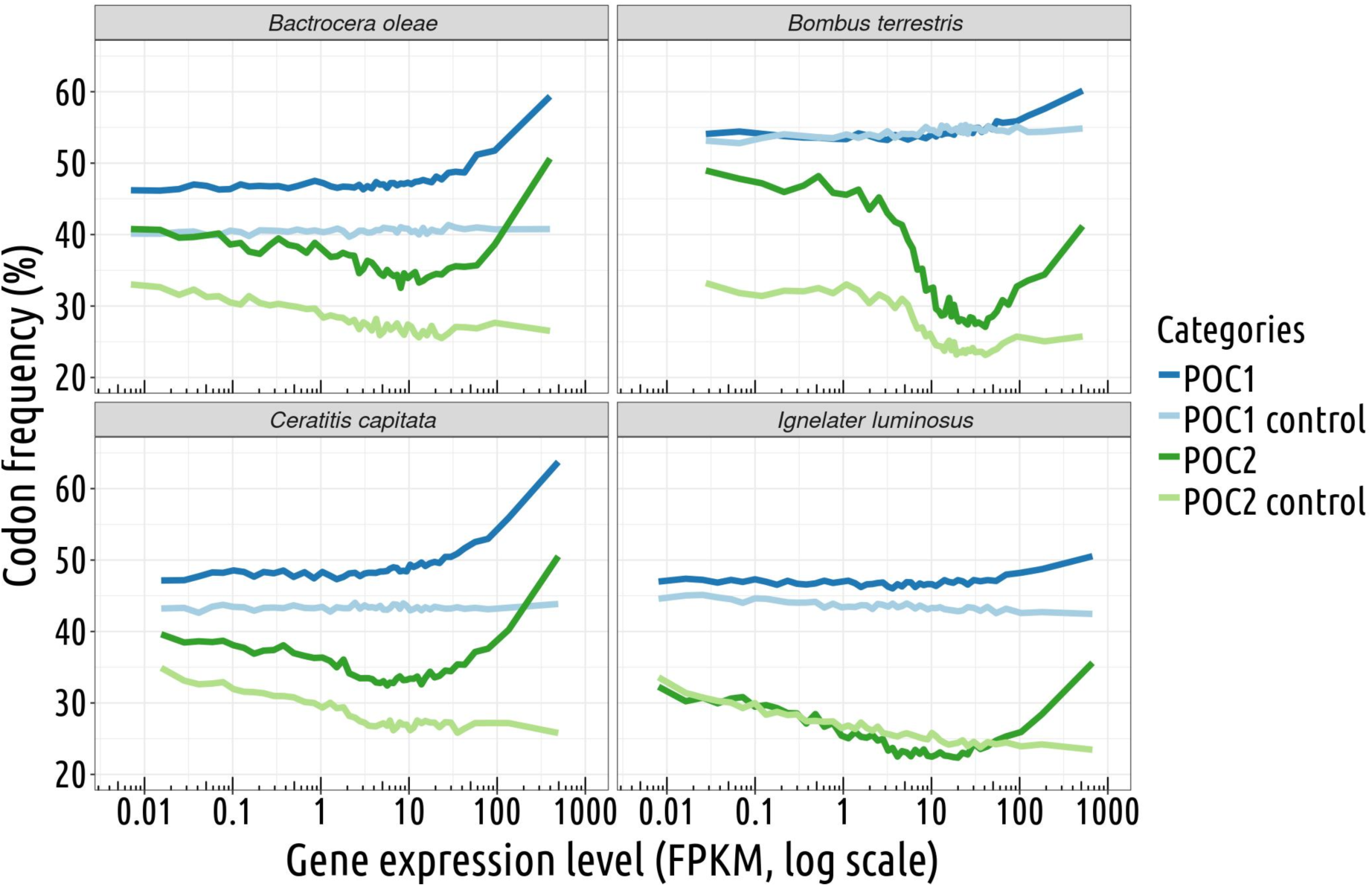
Relationship between the frequency of putative-optimal codons and gene expression level: examples of species showing variation in intron base composition with gene expression. Variation in the proportion of putative-optimal codons within coding sequences (POC1: dark blue; POC2: dark green) according to gene expression level. Variations in neutral substitution patterns, measured as the frequency of corresponding triplets within introns (POC1 control: light blue; POC2 control: light green). Each point represents a 2% bin of genes.

## Notes

https://zenodo.org/doi/10.5281/zenodo.12669922

### Competing Interest Statement

The authors have declared no competing interest.

